# Phase-separating fusion proteins drive cancer by dysregulating transcription through ectopic condensates

**DOI:** 10.1101/2023.09.20.558425

**Authors:** Nazanin Farahi, Tamas Lazar, Peter Tompa, Bálint Mészáros, Rita Pancsa

**Author notes:** Co-first authors (Nazanin Farahi, Tamas Lazar). Correspondence may be addressed to: Rita Pancsa or Bálint Mészáros.

## Abstract

Numerous cellular processes rely on biomolecular condensates formed through liquid-liquid phase separation (LLPS), thus, perturbations of LLPS underlie various diseases. We found that proteins initiating LLPS are frequently implicated in somatic cancers, even surpassing their involvement in neurodegeneration. Cancer-associated LLPS scaffolds are connected to all cancer hallmarks and tend to be oncogenes with dominant genetic effects lacking therapeutic options. Since most of them act as oncogenic fusion proteins (OFPs), we undertook a systematic analysis of cancer driver OFPs by assessing their module-level molecular functions. We identified both known and novel combinations of molecular functions that are specific to OFPs and thus have a high potential for driving tumorigenesis. Protein regions driving condensate formation show an increased association with DNA- or chromatin-binding domains of transcription regulators within OFPs, indicating a common molecular mechanism underlying several soft tissue sarcomas and hematologic malignancies where phase-separation-prone OFPs form abnormal, ectopic condensates along the DNA, and thereby dysregulate gene expression programs.

## Introduction

Many proteins and nucleic acids are able to undergo liquid-liquid phase separation (LLPS) and form biomolecular condensates in living cells^1^. These condensates, also frequently referred to as membraneless organelles (MLOs), are non-stoichiometric assemblies of macromolecules comprising a distinct liquid-like phase^2^ dedicated to specific cellular functions^3,4^. In the last few years, LLPS has emerged as a general and fundamental organizing principle employed by both prokaryotic and eukaryotic cells for the spatiotemporal segregation of their metabolic and signaling processes^5,6^.

Proteins play distinct roles in LLPS, classified as *scaffolds* (also termed as *LLPS drivers* but here we will reserve this word for cancer drivers), regulators and clients. *Scaffolds* can phase-separate on their own or in combination with other scaffolds (proteins, DNA or RNA), under native-like conditions. Regulators influence LLPS through affecting the expression, localization or modification states of the scaffolds. C*lients* do not influence LLPS but enter the condensates and may contribute to their functions^3,7^.

Although LLPS processes show a great heterogeneity in terms of the participating macromolecules and underlying molecular driving forces, they uniformly rely on multivalent weak/transient interactions between the (co)scaffolds that provide the flexibility required for the dynamic rearrangements crucial for LLPS^1^. Intrinsically disordered regions (IDRs), often of low sequence complexity, can play key roles in LLPS, usually by mediating weak residue-residue interactions^8–10^, or by carrying short linear motifs (SLiMs^11^) that bind to folded domains^12,13^. Homo-oligomerization is also frequently exploited by LLPS scaffolds to increase their valences^3,10,14^, and the binding of nucleic acids through RNA-or DNA-binding domains, or IDRs is also typical^15,16^. Elucidating the mechanism of formation, functions and regulation of LLPS systems remains a challenging task^17^. Nonetheless, many such systems have already been described, and several dedicated LLPS databases became available^5,18–20^ providing rich annotations enabling potential generalizations on the associated proteins^21^.

Numerous crucial cellular processes rely on phase-separated condensates, for instance, transcription and its regulation rely on RNA polymerase II condensates^22^, super enhancers^23^ and chromatin compartments with distinct histone modification patterns^24,25^. Therefore, perturbations of LLPS and the associated condensates can readily lead to the development of various diseases^26,27^. Phase-separated liquid-like structures can make a transition into less dynamic hydrogels or amyloid-like protein aggregates that are associated with certain neurodegenerative diseases^28,29^, such as amyotrophic lateral sclerosis^30,31^ and Alzheimer’s disease^32^. RNA-binding proteins are abundantly represented among LLPS scaffolds^10^ and are implicated in diverse diseases, such as neurodegenerative disorders, muscular atrophies and cancer^33^. The development of somatic cancers was generally attributed to the accumulation of driver mutations that alter the stability, activity or interactions of key proteins. However, recently it became evident that mutations can also interfere with and/or over-activate the formation of phase-separated condensates^34^. The presence or absence of certain MLOs are accepted diagnostic markers of certain cancer types^34^. For example, enlarged nucleoli are characteristic of large-cell lung carcinoma, or the lack of promyelocytic leukemia (PML) bodies are distinctive of acute promyelocytic leukemia^34^. Cancer-associated mutations directly affecting LLPS have only been demonstrated for some proteins^35–40^. Large-scale computational analyses highlighted that proteins implicated in diseases, including cancer, are enriched in predicted LLPS propensity^41^. Also, thousands of disease mutations have been identified in predicted LLPS scaffolds that likely contribute to condensate dysregulation^42^.

The role of LLPS in cancer has been extensively investigated and reviewed recently^25,43–49^. In two recent studies, cancer was linked with LLPS of the products of oncogenic fusions resulting from chromosomal rearrangements. In one, experimental and computational analysis of a large number of cancer-related fusion proteins have shown their propensity to be localized in cellular condensates^50^. In the other^51^, LLPS of a few fusion proteins combining phase-separating and DNA-binding regions have been experimentally confirmed. More interestingly, they have been shown to be targetable by small-molecule inhibitors.

Here, we take a rational approach to dissect these relations. By focusing on experimentally proven LLPS scaffolds and cancer drivers, we provide an unbiased assessment of the relevance of LLPS proteins in cancer compared to various other disease classes. We offer a multi-level description of the biological processes and functions that are preferentially associated with LLPS proteins involved in cancer and assess the underlying mutational mechanisms, showing the preponderance of fusion events, especially in certain early-onset somatic tumors. Using protein-region centric functional annotations, we show how they combine cellular functions with the ability to drive condensation, and how these newly emerging combinations of functional elements may offer novel ways of targeting this so-far largely undruggable class of oncogenes.

## Results

### 1. LLPS scaffolds play a significant role in the development of somatic cancer

In order to investigate potential links between liquid-liquid phase separation (LLPS) and proteins implicated in different groups of disease, we conducted a comprehensive analysis of the underlying associations. First, we tested how much LLPS scaffolds (**Table S1**) overlap with proteins involved in various classes of human diseases. We obtained four sets of proteins (see **Figure 1A-D** and **Table S2**) implicated in germline cancer, somatic cancer, neurodegenerative diseases and other human diseases (*see Data and methods*). For each of them, we generated 1000 random sets of human proteins with the same distributions of subcellular localizations and level of annotations (*see Data and methods and **Table S3***), and calculated the overlap with LLPS scaffolds. The distributions of overlaps between LLPS scaffolds and these random protein sets were compared to the true overlaps between LLPS scaffolds and the real disease-linked proteins of the four disease classes (**Figure 1A-D)**. Germline cancer proteins show an overlap with LLPS scaffolds that is indistinguishable from a random overlap. Other human diseases are slightly enriched in LLPS scaffolds, while neurodegenerative diseases are very significantly enriched with the observed overlap exceeding 6 standard deviations above the value expected at random. This finding conforms to previous studies elucidating the role of LLPS in neurodegenerative diseases^29,31^. However, LLPS scaffolds exhibit an even higher enrichment in somatic cancer driver genes, with the observed overlap being over 8 standard deviations higher than expected. Calculations done on a larger, but less confident dataset of LLPS scaffolds derived from PhaSepDB (**Table S4**) used as an independent alternative of our scaffold dataset also confirmed the observed tendencies (**Figure S1**). This shows that biological condensation is central to the development of somatic cancer in general. Proteins that regulate LLPS or partake in condensation in a client role only have a more moderate overlap with around 3 standard deviations above expectation level (**Figure S2**). Also, this effect is specific to condensation through LLPS, as proteins prone to aggregate through amyloid formation (**Table S5**) do not show a significant overlap with somatic cancer (**Figure S3**).

**Figure 1:**
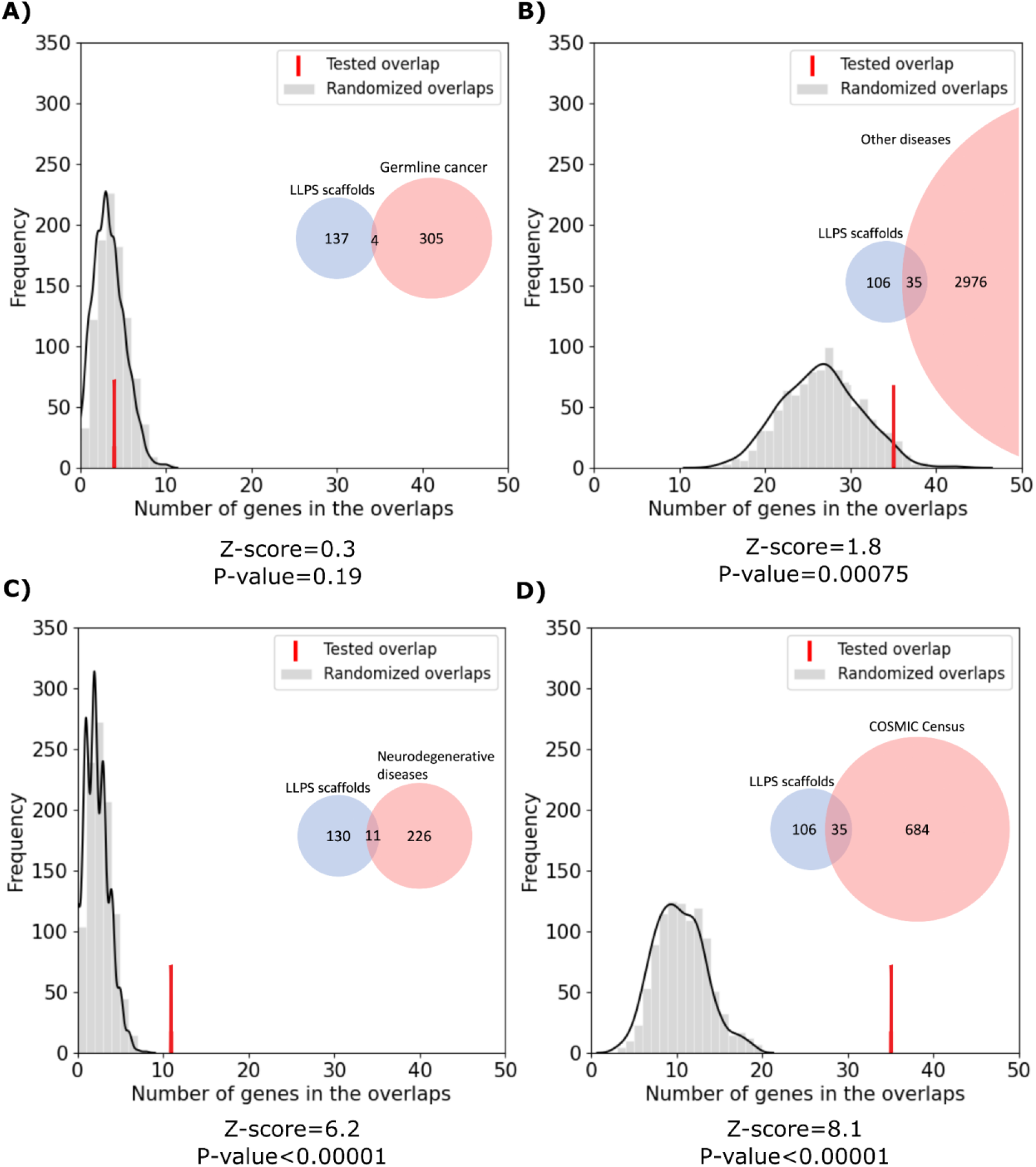
Overlap between LLPS scaffolds and various disease-associated proteins. Gray distributions show the expected overlap between LLPS scaffolds and the four classes of disease-associated proteins: (**A**) germline cancer, (**B**) other human diseases, (**C**) neurodegenerative diseases and (**D**) somatic cancer. Distributions were calculated from 1000 random generated sets of human proteins with subcellular localizations and levels of annotation matched to the real disease protein sets (*see Data and methods*). Red bars mark the observed overlap between the true protein sets with the corresponding Z-scores and p-values indicated below the graphs. Inset Venn diagrams show the number of proteins in each set and the observed overlap between them with circle areas being proportionate to the corresponding set sizes.

### 2. LLPS scaffolds are heavily associated with most cancer hallmarks

Tumor cells are known to acquire ten common phenotypes that are referred to as the cancer hallmarks^52^. Using annotations for the known cancer drivers in COSMIC Census (**Table S6**), we analyzed how often LLPS-related cancer proteins are connected to each of these hallmarks. We focused on LLPS scaffolds, regulators and clients that are annotated as cancer drivers in COSMIC and compared their involvement in each hallmark with those of all cancer drivers (**Table S7**). **Figure 2A** shows that LLPS scaffolds are enriched in most hallmarks, with statistical significance (p<10e-2) in seven. PhaSepDB-derived scaffolds also confirmed this tendency (**Figure S4**). LLPS regulators also exhibit significant enrichments in four hallmarks, while LLPS clients show no significant enrichment. **Figure 2B** also shows that LLPS scaffolds more often have an effect on several hallmarks than cancer drivers in general. While hallmark annotations are obviously sparse (roughly half of all cancer drivers are not associated with any hallmark), there is a clear tendency for LLPS scaffolds influencing several hallmarks. Over 25% of cancer-driving LLPS scaffolds contribute to 5 or more hallmarks, while in general this is only true for less than 10% of all cancer drivers.

**Figure 2:**
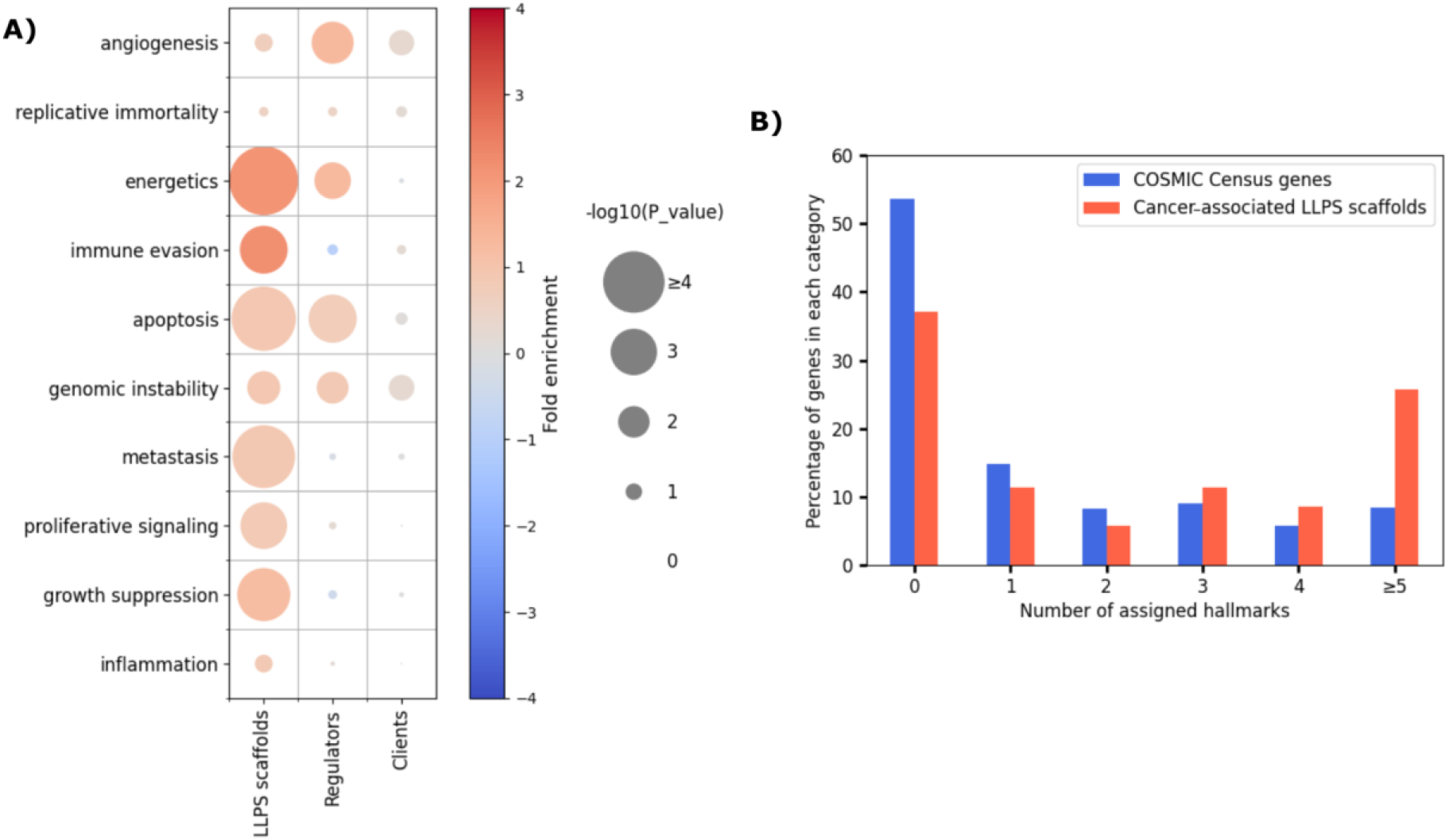
Association of LLPS-related proteins with cancer hallmarks. (**A**) The color of the circles in the heatmap represents the fold enrichment, while the size represents the significance of overrepresentation/depletion for the three classes of LLPS-related proteins in the ten hallmarks of cancer. (**B**) The histogram depicts the number of hallmarks individual cancer driving LLPS scaffolds contribute to (red) as compared to cancer drivers in general (blue).

### 3. Cancer-associated LLPS proteins are enriched in critical molecular functions, including mRNA processing, transcription regulation and chromatin remodeling

To evaluate the molecular mechanisms of LLPS-related proteins in cancer, we defined ‘molecular toolkits’, sets of Gene Ontology terms that capture a high-level molecular function (*see **Table S8** for toolkit definitions*). We grouped our toolkits into 5 supertoolkits that cover broad, cell-level functions (**Table S9**).

Molecular toolkits evaluated for cancer drivers show that the most commonly affected functions are heterochromatin organization, DNA binding and gene silencing, protein maturation and stability, cell surface receptor signaling, intracellular signal transduction, intracellular transport, and cell adhesion (**Figure 3**). Compared to cancer drivers’ toolkit enrichments, LLPS scaffolds, regulators and clients all have distinct toolkit repertoires (**Table S10**). Cancer driver LLPS scaffolds are most significantly linked to mRNA processing, its regulation, and chromatin remodeling (see **Figure 3B** for a selection of toolkit terms that show significant enrichments in LLPS-related cancer drivers). Dysregulation of LLPS regulators impacts a lot more molecular processes including DNA recombination and replication, protein localization to organelles, translation, regulation of gene silencing and epigenetic regulation of gene expression (**Figure 3B**). Finally, molecular functions of LLPS clients altered by cancer are most commonly associated with chromosome segregation, actin and microtubule cytoskeleton organization and with regulation of substrate adhesion-dependent cell spreading (**Figure 3B**).

**Figure 3:**
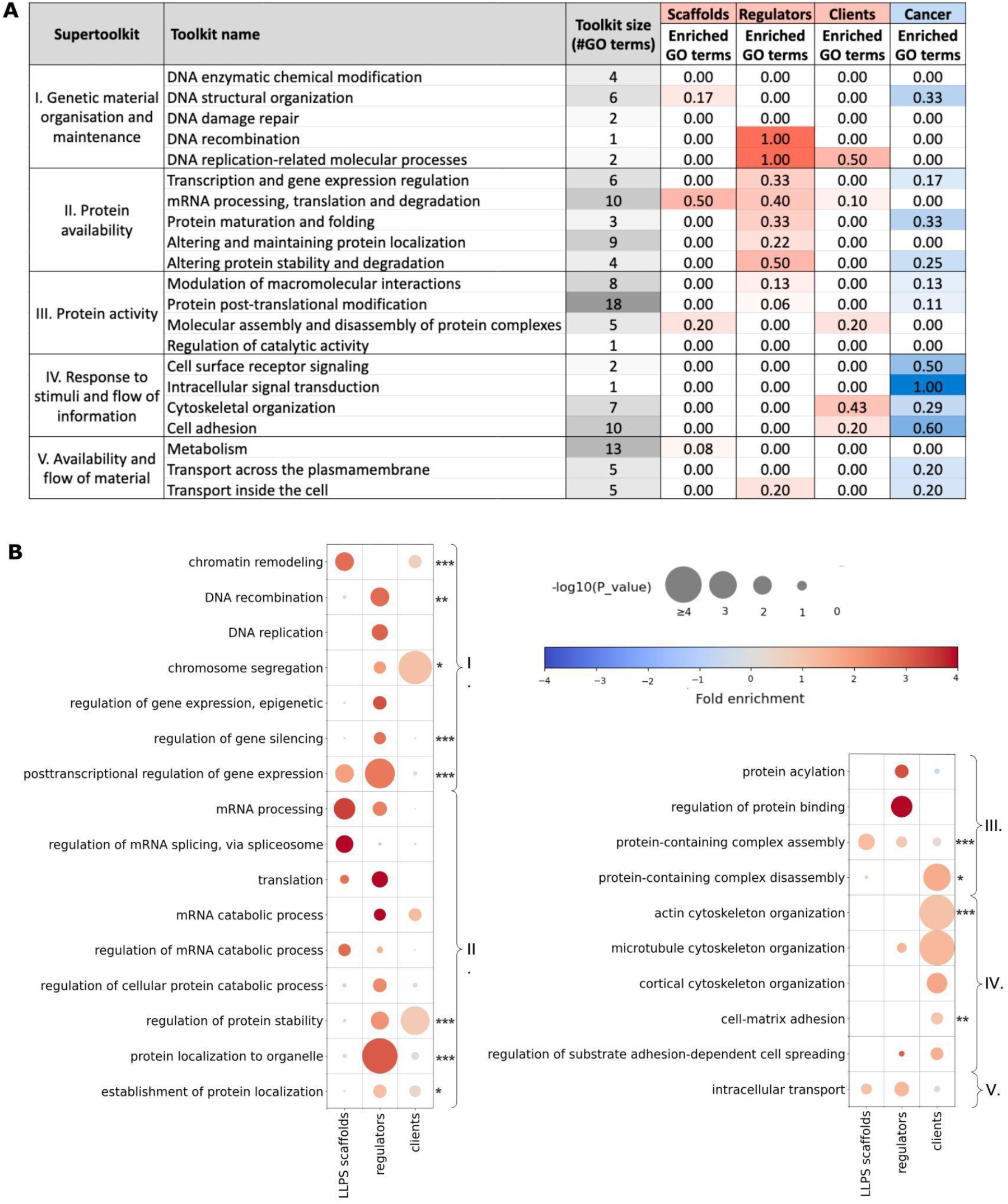
Enrichment of functional toolkits in the three classes of LLPS-related proteins (scaffolds, regulators, clients). (**A**) A particular toolkit was considered to be enriched in a protein class if the fold enrichment was ≥1 and p-value of significance was <0.05 in Fisher’s exact test. (**B**) The heatmaps depict significantly enriched GO terms (fold enrichment > 1.0 by Fisher’s exact test) with a minimum of 3 proteins in a given LLPS class. On the right side of the heatmap, one, two or three stars indicate the significance level of the GO enrichments for the whole set of cancer drivers compared to a random background (with levels 0.05>p≥0.01, 0.01>p≥0.001, p<0.001, respectively). GO terms belonging to the same supertoolkit are connected by brackets with the numbering of supertoolkits also provided. Definitions of toolkits by GO terms and their unfiltered individual fold enrichment and significance values are listed in **Tables S8 and S10**.

### 4. Cancer-associated LLPS scaffolds typically drive cancer via dominant gene fusions and lack available drugs

To better understand their roles played in cancer development, we tested various features of cancer-associated LLPS scaffolds, regulators and clients in comparison to all known cancer drivers from COSMIC Census (**Table S6**). **Figure 4A** shows that LLPS scaffolds are enriched in oncogenes (see also **Figure S5** where enrichment in oncogenes and tumor suppressors is confirmed based on comparisons against equivalent randomized background sets) and are preferentially affected by dominant mutations (**Table S7**). In contrast, LLPS regulators are slightly enriched in tumor suppressors and are targeted by dominant mutations to a much lower degree. LLPS clients show no enrichment in either tumor suppressor/oncogene role or in dominant/recessive mutations. This shows that on average the more significant role a protein plays in phase separation (with scaffolds > regulators > clients), the more likely it is to be an oncogene affected by dominant mutations.

**Figure 4:**
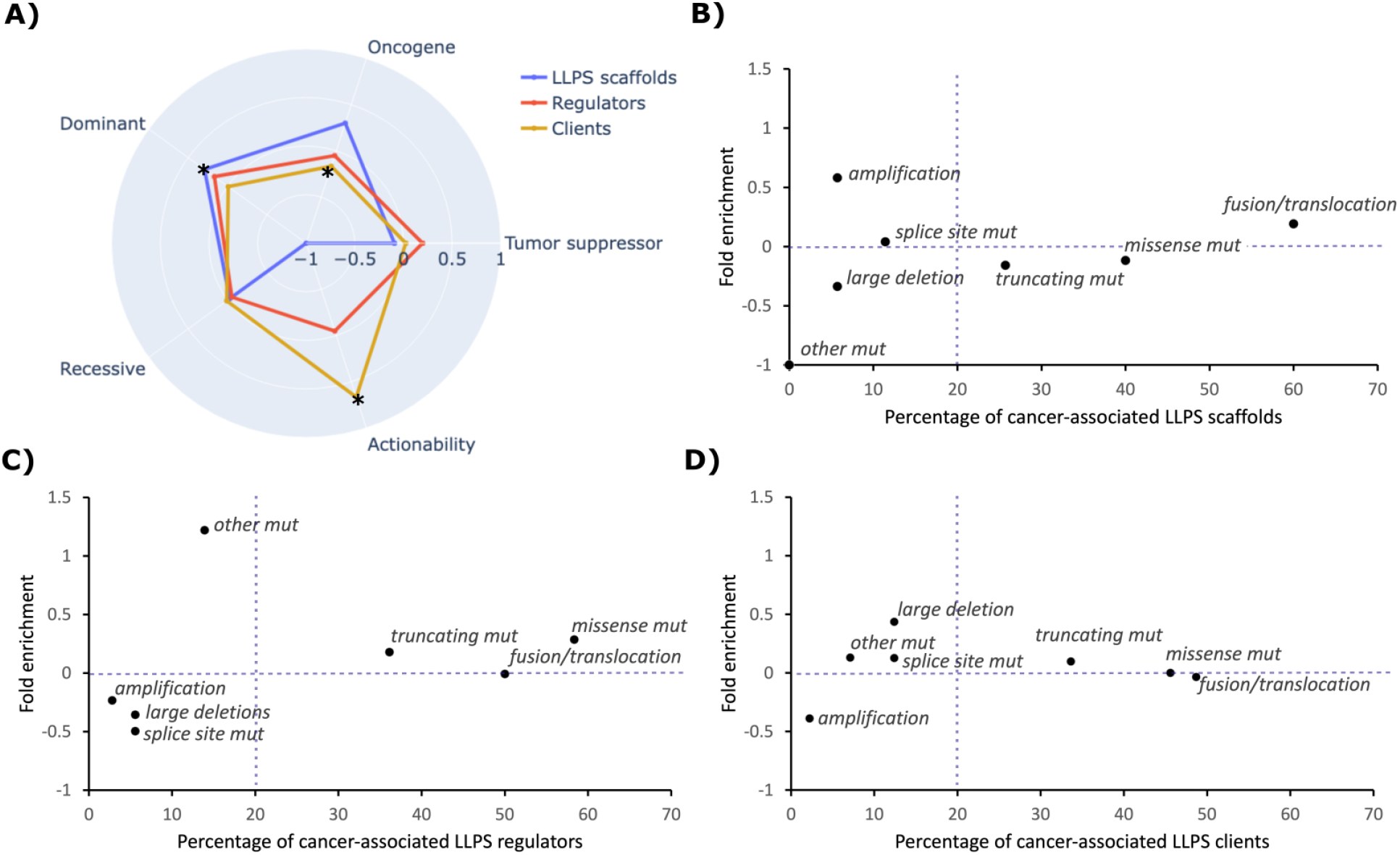
Characteristic features of LLPS-related cancer driver proteins compared to cancer drivers in general. (**A**) The radar chart plots the fold enrichments of the three classes of cancer-associated LLPS proteins in cancer drivers that are oncogenes or tumor suppressors; that are affected by dominant or recessive mutations; and those with available FDA-approved drugs (actionability). (**B-D**) The percentage of cancer-associated LLPS scaffolds (**B**), regulators (**C**) and clients (**D**) affected by various dominant mutation types is presented on the x axis, while their fold enrichment values compared to COSMIC census as background is presented on the y axis. The truncating mutation category comprises both frameshift and nonsense mutations.

Comparing the actionability – i.e. the number of FDA-approved drugs available – for various proteins, LLPS scaffolds show an extreme depletion (**Figure 4A**). None of the 35 cancer-associated scaffolds have any FDA-approved drugs, even considering off-label standard care use (*see Data and methods for definitions*). LLPS regulators involved in tumorigenesis are also relatively poorly targetable, since only 2 out of the 36 proteins – KRAS and BRCA1 – have available drugs that act on them. In contrast, LLPS clients are generally the most actionable with 22 out of 226 having available drugs (**Figure 4A** and **Table S7**).

Analyzing specific types of genetic alterations shows that all three classes of LLPS proteins are mostly affected by the same three mutation types (**Figure 4B-D**): missense mutations (which have a local effect on the protein); frameshift and nonsense mutations (which truncate the protein); and translocations/fusions (which can create new proteins by combining regions of independent proteins into a single product). Fusions/translocations were found to be the most abundant mutation type for LLPS scaffolds. Although being only slightly enriched in this mutation type compared to cancer drivers in general, 60% of the LLPS scaffolds form oncogenic fusions and for most of them this is the sole mutation type observed in cancers (**Figure 4B**). In contrast, LLPS regulators are enriched in and are most often affected by missense mutations, while LLPS clients show no enrichment in missense mutations and fusions/translocations compared to cancer drivers in general. The tendencies proposed for LLPS scaffolds are mostly confirmed by calculations made on PhaSepDB-derived scaffolds (**Figure S6**).

### 5. Oncogenic fusion proteins represent novel combinations of functions driving tumorigenesis

The analysis of COSMIC annotations clearly highlighted that LLPS scaffolds primarily contribute to cancer through forming oncogenic fusion proteins (OFPs). Therefore, we performed a systematic analysis of the known OFPs of COSMIC Census proteins. When assembling the OFP dataset, we deliberately followed a highly selective approach in order to exclude passenger OFPs, which were just once observed in patient samples through sequencing approaches and do not necessarily play a causal role in the respective cancer type. For this reason we only used COSMIC-curated fusions and chose not to take OFPs from the TCGA database (www.cancer.gov/tcga/), unlike the two recent studies analyzing the predicted LLPS propensities of OFPs^50,51^, who optimized on the abundance of data. Since the vast majority of the 32 TCGA cancer types are typically not primarily relying on gene fusions/OFPs, while many rare cancer types are, we turned to COSMIC, where rare cancer types often defined by the presence of a well-defined OFP (or a group of those) are also covered, and only well-documented fusions of the Census proteins are listed that recurrently occur in a particular cancer type and have a widely recognized role in driving oncogenesis. Of the 450 unique fusion gene pairs identified for COSMIC Census genes, 303 in-frame-fused chimeric OFPs could be obtained wherein each gene pair is represented by a single OFP and the fusion boundaries could be precisely defined on the protein level (**Table S11**). Due to different data selection strategy and the inclusion of rare cancer types, our 303 COSMIC-derived cancer driver OFPs only show a limited overlap with the large sets of fusion oncoproteins recently analyzed for predicted LLPS propensities by Wang *et al.*^51^ and Tripathi *et al.*^50^ (**Figure S7A**), therefore most of our annotated fusions remain unique to our dataset and have never been investigated in relation to LLPS.

We analyzed our OFPs by scanning them for known conserved protein modules using Pfam, InterPro and UniProt annotations (*see Data and methods and* **Tables S12-14**). Since the fusion breakpoints of well-characterized OFPs tend to reside in disordered protein regions, leaving folded domains intact^53^, we did not have to deal with domains/modules cut into half by the fusions. Many protein modules perform similar functions, therefore, we aimed at analyzing the molecular functions conveyed by them. These functions can be captured using Gene Ontology (GO) terms, however, GO terms are assigned to full proteins. To enable a systematic analysis of the associations between functions in the fusions, GO molecular functions and biological processes were assigned to the protein modules of the collected OFPs and their wild-type constituent proteins (**Figure 5A**; **Tables S15**-**S16**; see *Data and methods* for more details). The GO terms assigned to the protein modules were mapped to a GO subset (GO Slim) representing biologically relevant, fairly specific, yet high level processes and functions (**Table S17**). Pairwise association levels were then determined for each possible pair of these processes/functions assigned to the modules of the wild-type constituent proteins or OFPs, separately (see **Tables S18-S19** for the resulting functional association matrices). The functions exhibiting significantly higher association levels in OFPs were considered to be fusion-specific (**Figure 5A-B; Table S20**).

**Figure 5:**
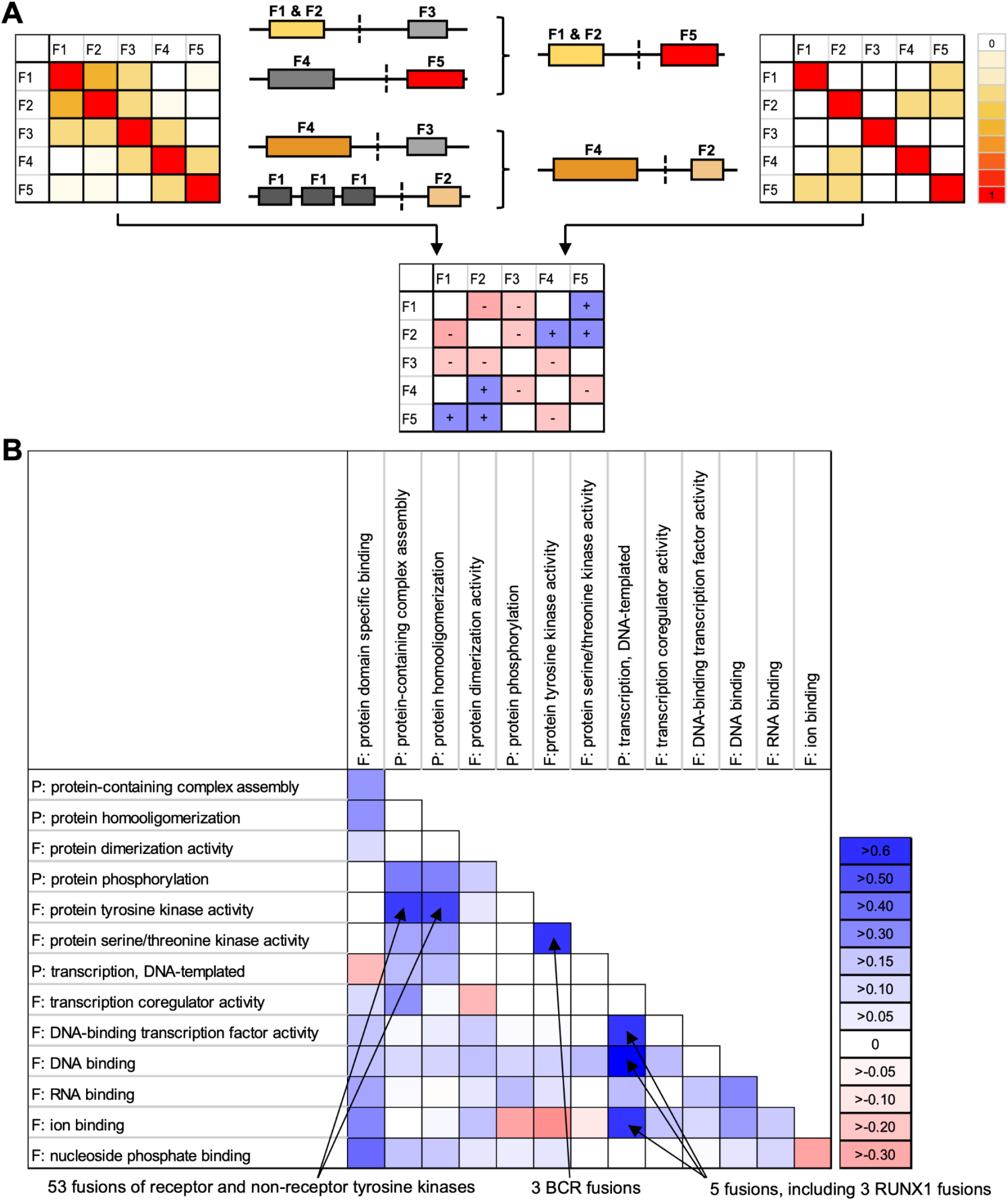
Oncogenic fusion proteins represent novel combinations of molecular functions/processes. (**A**) The functional modules of oncogenic fusion proteins (OFPs) and wild type constituent proteins were annotated by module-specific molecular functions/processes. Pairwise association levels between the annotated functions were calculated using overlap coefficients for OFPs and their constituent proteins separately. The association levels calculated for constituent proteins were subtracted from those calculated for OFPs to highlight the function pairs whose association levels are increased or decreased in OFPs. (**B**) Heatmap showing the pairwise associations between functions (based on our GO Slim definition provided in **Table S17**) after removal of redundancy, for those terms that are increased (shades of blue) or decreased (shades of red) in OFPs considerably (ΔOC ≥ 0.4 in **Table S20**). Three additional terms are shown that do not fulfill the previous criterion but are important for our study and generally considered to be linked to biomolecular condensation: protein phosphorylation, transcription coregulator activity and RNA binding. The associations that showed the largest increase in OFPs (ΔOC > 0.5) are labeled with the related fusion types. When calculating overlap coefficients, the number of elements in the intersection of the two sets is divided by the size of the smaller set, therefore the resulting value will range from zero to one, irrespective of the sizes of the two sets. Thus, it is possible for a set with only 3 BCR fusions to yield a comparable OC value as over 50 RTK fusions.

Our data highlight strong fusion-specific association between tyrosine kinase activity (and the associated protein phosphorylation function) and protein homooligomerization. Although the fusions of different receptor tyrosine kinases (RTKs) are implicated in different cancer types, they all rely on very similar molecular principles. In such fusions, RTKs lose their N-terminally encoded extracellular ligand-binding domains and commonly their transmembrane segments too, while the fusion partners replacing those can form homodimers or homo-oligomers. This leads to the dimerization, cross-phosphorylation and constitutive activation of the tyrosine kinase domains and relocalization to the cytoplasm or nucleus (depending on the partner). Consequently, this pathogenic process results in uncontrolled, ligand-independent phosphorylation of their downstream target proteins^54,55^. Although this association has been long recognized, to our knowledge, it has never been confirmed systematically on statistical terms. Our dataset reveals over 50 OFPs showing an association of these two functions (an overlap coefficient (OC) of 0.62 on a 0 - 1 scale for OFPs) while it does not occur in wild-type proteins (OC is 0.06; ΔOC=0.55) (**Figure 5B, Tables S18, S19 and S20**). Breakpoint cluster region protein (BCR) fusions important in chronic myeloid leukemia combine protein tyrosine kinase activity with protein serine-threonine kinase activity, which does not exist in wild-type proteins (ΔOC=0.6). However, these fusions also represent a subset of the fusions that combine oligomerization with tyrosine kinase activity. Oligomerization through BCR and compromised regulation of the fused non-receptor tyrosine kinases lead to their over-activation, which is central to oncogenicity (similarly to RTK fusions)^56–58^. Coupling of domains implicated in transcription directly or as activators/repressors (covered by the term “transcription, DNA templated”) with DNA-binding domains (ΔOC=0.8 for “DNA-binding”), mainly zinc finger (ZnF) domains (ΔOC=0.6 for “ion-binding”) of transcription factors (ΔOC=0.6 for “DNA-binding transcription factor activity”) is also specific for certain fusions, e.g. a subset of RUNX1 fusions. In the OFPs of RUNX1/AML1 fused to members of the CBFA2T family, the N-terminal DNA-binding RUNT domain of RUNX1 gets coupled to the TAFH, NHR2 and MYND domains of CBFA2T family proteins, of which the TAFH domain is a protein-binding module involved in transcription regulation^59^. A similar coupling is seen in the CBFA2T3-GLIS2 fusion where the TAFH domain of CBFA2T3 gets coupled to the DNA-binding C2H2-type ZnF domains of GLIS2, and also in the KMT2A-ELL fusion, where the DNA-binding CXXC-type ZnF of KMT2A gets coupled to ELL, which is part of the transcription elongation factor complex, thus having a direct role in transcription. In all, our results indicate that OFPs often exert their oncogenic effects through highly specific combinations of molecular functions and that our data and approach are well-suited to detect those.

### 6. Most oncogenic fusions of LLPS scaffolds couple phase separation with DNA-binding

Encouraged by the robust detection of function combinations already known to drive cancer, we introduced “driving biomolecular condensation” as a novel molecular function term and assigned it to the regions of LLPS scaffolds proposed to be minimally required for driving LLPS (**Table S21**). Then we determined if the novel term shows any fusion-specific associations to other functions/processes as described in the previous section (**Figure 6A**). Notably, most of the LLPS-driver regions of scaffolds are largely retained in their fusions (in 69 of the 72 fusions), thus they were assigned with the “driving biomolecular condensation” term. So, at least 69 fusions are expected to form condensates through LLPS due to inheriting LLPS-driver regions. For 14 of these 69 fusions the ability to drive phase separation is supported by two recent studies, where condensate localization of altogether 124 fusion oncoproteins has been demonstrated in cells^50,51^, while it was not disproved for any of them (**Figure S7B**). Interestingly, LLPS scaffolds tend to be located on the N-terminus of the fusion products (in 63 of 72 cases; **Figure 6B**), and since fusions always inherit the promoter and other gene regulatory regions of the N-terminally fused gene, their expression will be mostly regulated by the gene regulatory regions of the LLPS scaffolds. These LLPS-prone OFPs are typically implicated in early-onset soft tissue sarcomas and hematological malignancies, while they are less involved in the development of the otherwise abundantly occurring brain tumors, as well as late-onset carcinomas and skin cancers (**Figure 7**).

**Figure 6.**
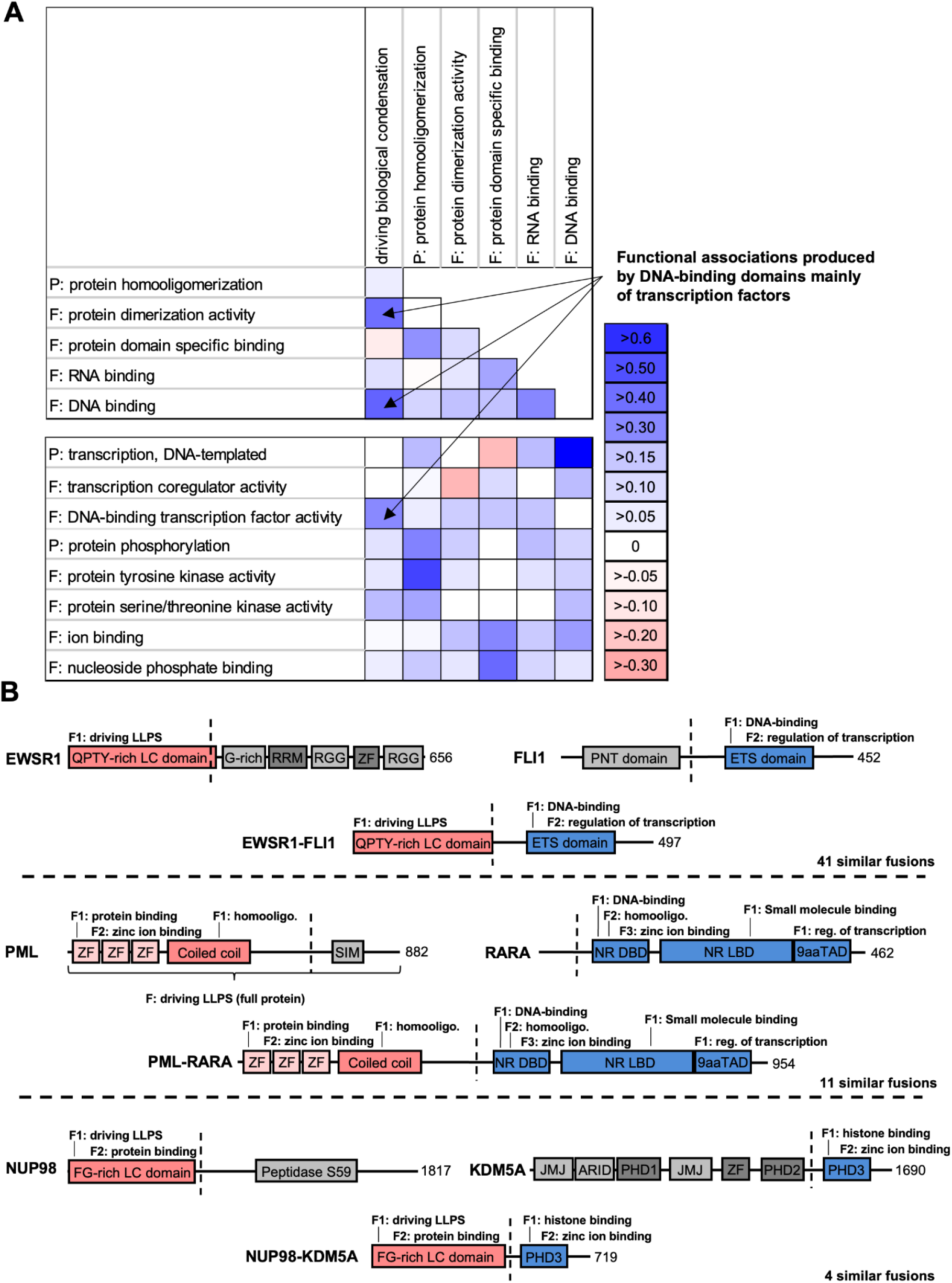
Functional associations in the fusions of LLPS scaffolds. (**A**) The previous module-level functional annotations were complemented by assigning the function “driving biomolecular condensation” to the regions of LLPS scaffolds that are minimally required for LLPS. Pairwise association levels were calculated between “driving biomolecular condensation” and other molecular functions important in condensate formation that capture the increase in valency or interaction capacity of the protein (see symmetric heatmap on the top) as described in Figure 5, as well as between the condensate formation-associated and the previously analyzed (Figure 5) molecular functions (‘F’) and biological processes (‘P’) (asymmetric heatmap in the bottom). The heat maps show how pairwise associations between functions change in OFPs (shades of blue for increase, shades of red for decrease). The functions associated with “driving biomolecular condensation” that showed the largest increase in OFPs (ΔOC > 0.2) are labeled. (**B**) Domain maps and assigned module-level functions of three well-studied, representative OFPs and their constituent proteins. Breakpoints of translocation are indicated with vertical dashed lines. Assigned functions are only depicted for the protein modules that are retained in the fusions, others are colored in shades of gray. Domains colored in red drive LLPS or homooligomerization, domains colored in blue mediate transcription by DNA-binding or chromatin(histone)-binding. The different zinc fingers (ZFs) of PML are colored light pink because they are not known to crucially contribute to the oncogenesis of the fusion protein. The sizes of proteins and domains are not proportionate to each other. LC - low complexity, RRM - RNA-recognition motif, ETS - erythroblast transformation specific domain, PNT - pointed domain, SIM - SUMO-interacting motif, NR - nuclear receptor, DBD - DNA-binding domain, LBD - ligand-binding domain, TAD - transactivation domain, PHD - plant homeodomain, JMJ - jumonji domain, homooligo. - homooligomerization, reg. - regulation.

**Figure 7:**
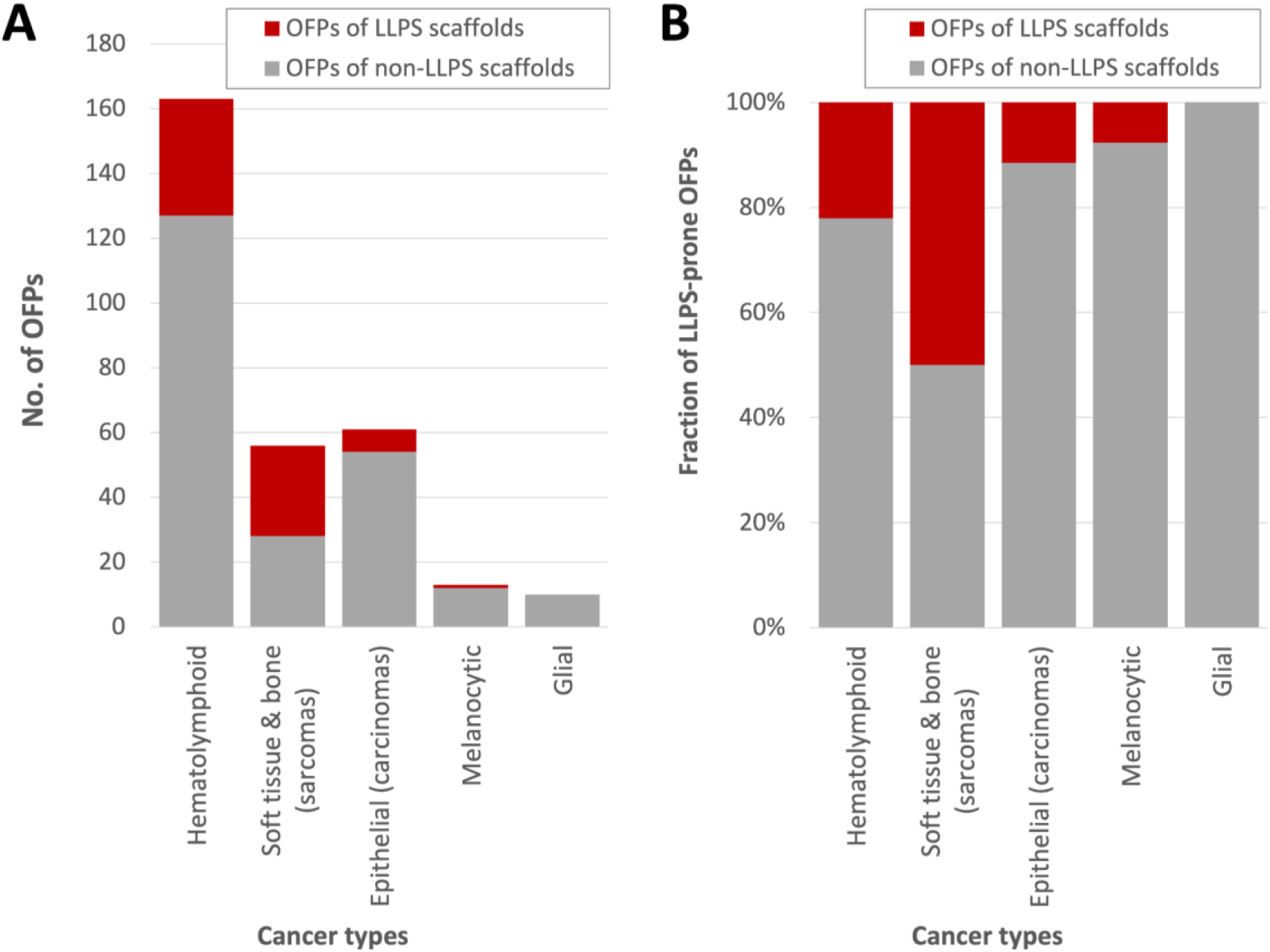
The fraction of LLPS driver region-containing oncogenic fusion proteins (OFPs) in different categories of cancers. The absolute number (**panel A**) and relative percentage (**panel B**) of OFPs are shown in the five big categories of cancers defined by the developmental origin of the cancerous tissue. The OFPs that contain a known LLPS driver region are colored red, while those that do not are colored gray. For each OFP, the respective cancer type was obtained from the original data source (COSMIC/UniProt/original article). In the rare cases when more than one cancer type was indicated for the same OFP, the most frequently associated one was selected. Then these cancer types were grouped into five major categories defined by the developmental origin of the cancerous tissue (Supplementary Table S11). OFPs within the category of hematolymphoid cancers are implicated in lymphomas, leukemias and other neoplasms of myelocytes (e.g. myelodysplastic syndromes). OFPs of the soft tissue and bone category underlie the development of sarcomas but also some benign tumors of the soft tissues, such as lipomas. The OFPs observed in epithelial cancers (i.e. carcinomas) make up a distinct category. The category of melanocytic cancers involves OFPs implicated in skin tumors, such as melanomas and Spitz tumors. The category of glial cancers encompasses OFPs observed in malignancies of glial cells, including astrocytes.

The molecular function “driving biomolecular condensation” showed the strongest fusion-specific increase in association levels with the functions “DNA binding” (ΔOC=0.46), “protein dimerization activity” (ΔOC=0.42) and “DNA-binding transcription factor activity” (ΔOC=0.35) (**Figure 6A**). Other functions/processes did not exhibit strong (ΔOC>0.3) changes in associations. Association with “DNA binding” was found in 52 of the 69 LLPS-prone OFPs, a coupling of functions that has also been captured by Wang *et al.*^51^. Notably, “driving biomolecular condensation” and “DNA binding” are also moderately associated in wild-type proteins (OC=0.38), probably because many transcription factors (TFs) have been reported to phase-separate under certain conditions^60–62^. The other three detected functional associations were somewhat weaker and were identified in subsets of the DNA-binding fusions – this is not surprising, since a domain could have multiple annotated functions, and most DNA-binding domains occur in TFs, many of which dimerize.

In 41 of the 52 fusions (all EWSR1, FUS and TAF15 fusions and 12 NUP98 fusions) that are typically implicated in soft tissue sarcomas and hematological malignancies, respectively, a potent low-complexity LLPS-prone region is coupled with an intact DNA-binding domain of certain transcription factors (mainly ETS domain-containing TFs in FET fusions and homeobox TFs in NUP98 fusions) (**Figure 6B, Table S11**). In the other 11 fusions with the same association, including PML-RARA, NPM1-RARA, BCOR-RARA, NUMA1-RARA, STAT5B-RARA, ZBTB16-RARA, NPM1-MLF1, NONO-TFE3, SFPQ-TFE3, PAX5-ELN and PAX5-PML, which are mainly associated with acute leukemias, an oligomerization-prone subregion of an LLPS-driver or any other protein is combined with a TF (**Figure 6B**). (Since retinoic receptor alpha (RARα) is an LLPS scaffold and it combines LLPS-prone disordered regions with a DNA-binding domain in itself^60^, all its fusions are a part of the dataset.) The LLPS propensity of the TFs is likely increased in their oligomerization-prone fusions due to increased multivalency. At the same time, homo-oligomerization through an extraneous domain can compromise certain functional modalities of the incorporated TFs (as seen for PML-RARA^63,64^ and NONO-TFE3^65^). Interestingly, these fusions not only differ from the fusions of the previously introduced group based on the properties of the incorporated LLPS-prone regions, but they also show different pathomechanisms. Most of them were shown to exert a dominant negative effect on the transcriptional activity of the incorporated TFs that depends on the oligomerization of the fusion partners^64,66–69^. Also, they may recruit activating and repressing chromatin remodeling complexes to deregulate transcription^64,70^.

Manual inspection of the domain structures of the 17 fusions that did not combine “driving biomolecular condensation” with “DNA-binding” showed that 4 combine the LLPS-driver region of NUP98 with chromatin-binding domains (displaying a similar pathomechanism to DNA binding NUP98 fusions^71^) (**Figure 6B**), 5 show associations between different oligomerization-prone subregions of LLPS scaffolds and tyrosine kinase domains of RTKs (the molecular pathomechanism of these has been described in the previous section), while the remaining 8 represent unique (e.g. DNAJB1-PRKACA^72^) or not completely clear functional associations.

## Discussion

We set out to systematically study the connection between cancer and biological condensation, specifically mapping the extent to which LLPS is affected in cancer and understanding the molecular pathomechanisms and therapeutic consequences of mutations affecting LLPS scaffolds. Our motivation is driven by our observation that out of diseases with a known causative protein repertoire, somatic cancer has the strongest connection to LLPS scaffolds, far surpassing those of other diseases, including neurodegenerative disorders where several such LLPS scaffolds are linked to disease emergence (**Figure 1**). In contrast, germline cancer mutations are extremely rare in LLPS scaffolds, indicating that these mutations have a strong phenotypic effect, not tolerated to occur ubiquitously in the whole body. Our high-level disease grouping demonstrates that there might be a correlation between disease severity and involvement of LLPS, as many somatic cancers have much faster progression, if untreated, as compared to cancer predisposition syndromes arising from germline cancer mutations or compared to neurodegenerative diseases. This indicates that the modulation of LLPS scaffolds via cancer mutations produces strong phenotypes. We focused on various aspects of tumorigenesis, ranging from mutational mechanisms, through modulation of biological processes, up to the emergence of cellular hallmarks, to understand why and how this happens.

Our data show that cancer-driving LLPS scaffolds are potent oncogenes, giving rise to dominant phenotypes and lacking targeting options by current FDA-approved drugs (**Figure 4A**). These properties not only contrast LLPS scaffolds with cancer drivers in general, but also with cancer drivers playing a regulator or client role in LLPS. Therefore, the mutation or dysregulation of proteins directly involved in inducing biological condensation gives rise to the most detrimental phenotypes. Many studies have provided insights into these genetic alterations showing that overexpression or missense mutations can produce gain or loss of function for LLPS scaffolds^25,43–48^. However, we found that 60% of the cancer-driving LLPS scaffolds are predominantly affected by gene fusions that create oncogenic fusion proteins (OFPs) (**Figure 4B**). This is in agreement with individual cases where LLPS scaffolds were found to contribute to different cancer types through forming OFPs^48,73^. OFPs display diverse pathomechanisms^74^, they could alter the regulation or localization of important hub proteins thereby rewiring protein interaction networks^75–77^, and/or introduce specific combinations of protein domains/functions that have a high potential for driving cancer^47,53,54^. In a high-throughput study, a large set of TCGA-derived OFPs (with yet-unvalidated roles in the respective cancer types) were analyzed for various LLPS-associated predicted features and 166 were tested for punctate/condensate localization in HeLa cells^50^. This study concluded that the majority of fusion oncoproteins are likely to partition into condensates, and highlighted important physicochemical features associated with nuclear and cytoplasmic condensation. Furthermore, they derived 4 major archetypical classes of OFPs, and using the set of computed features developed a prediction tool to analyze the LLPS-propensity of OFPs in high throughput^50^.

Importantly, several OFPs of LLPS scaffolds have been already shown to undergo LLPS, such as those of the FET family proteins (<u>F</u>US, <u>E</u>WSR1 and <u>T</u>AF15)^78–82^ and nucleoporins^73,83–85^, and some others, such as NONO-TFE3^65^, SS18-SSX^86^, BRD4-NUTM1^87^, SFPQ-TFE3^51^. Most of these fusion products are primary drivers of cancer (primarily of sarcomas and hematolymphoid cancers, as shown in **Figure 7**), i.e. they are potent oncogenes with the ability to drive the tumorigenic transformation of healthy cells by themselves^64,88–92^. In their case, oncogenicity is mostly attributed to their ability to form condensates at non-native subcellular locations^48^.

The mechanism of action of OFPs fundamentally differs from other cancer-mutated proteins^74^, as they can combine molecular functions in a novel way that is detrimental to the healthy cell, driving oncogenic transformation^53^, as exemplified by RTK fusions^25,54,76^. We explored this functional association by attaching functional annotations to protein regions that can be identified in any protein sequence in an automated and high-throughput way (**Figure 5**). Through systematic analysis, we found that the vast majority of OFPs that contain regions of LLPS scaffolds inherit the ability to drive phase separation, and we propose that they can be classified into 4 main categories: low complexity LLPS scaffolds coupled with DNA-binding via 1) transcription factor (TF) domains or 2) chromatin-binding domains; and oligomerization-prone subregions of LLPS scaffolds fused to 3) TFs or 4) receptor tyrosine kinase domains (**Figure 6**). Category 1 is specific for soft tissue sarcomas (FET family fusions) or acute leukemias (NUP98 fusions), categories 2 and 3 are mainly responsible for acute leukemias, and category 4) shows no obvious cancer type specificity. Nonetheless, fusions of the first three categories all seem to rely on similar molecular principles, representing potent, LLPS-prone transcriptional activators^78^ or repressors^93^.

A likely reason for the strong detrimental phenotypic effect of LLPS-scaffold OFPs belonging to the first three categories is that the combination of TF activity with the ability to self-sufficiently initiate phase separation is uncommon in a healthy cell. Wild-type TFs tested for LLPS so far could only phase-separate on their own at high concentrations^60,61^, which is in conflict with their otherwise notoriously low cellular levels^7^. At near-physiological concentrations, TFs require at least a coactivator and a specific DNA segment for LLPS^62^, therefore, they are typically context- and partner-dependent LLPS scaffolds. In contrast, in the context of fusions TFs are complemented by potent LLPS-driver regions or at least by homo-oligomerization domains and display elevated expression levels due to the exchange of their gene regulatory regions, which both favor condensate formation. Therefore, such fusions resolve the dependencies of the incorporated TFs and form ectopic condensates along the DNA even at genes which are not normally regulated by the TF^69,94,95^. Such condensates act as potent transcriptional activators or repressors by efficiently recruiting diverse chromatin remodeling complexes^64,69–71,96–101^ (or even RNA polymerase II itself^78,102^), leading to aberrant gene expression patterns^34,103^. The pathomechanisms of many fusions in our dataset (see **Figure 6B** for examples and **Table S11** for a full list of fusion constructs) have been studied individually, however, our results underscore that they represent a much larger group of LLPS-prone OFPs that combine similar functions and thus likely rely on similar underlying molecular principles.

This unique molecular makeup of LLPS-scaffold OFPs is reflected in the biological processes the cancer-associated LLPS scaffolds are involved in. By performing enrichment analysis using standard Gene Ontology (GO) terms, we can recapitulate that the most affected processes are chromatin remodeling, as well as mRNA-related terms (**Figure 3**). For instance, the nucleolar protein nucleophosmin is a regulator of mRNA splicing that functions in chromatin remodeling^104–106^, however, it often forms oncogenic fusions resulting in loss of function in lymphomas^107^. In contrast, regulators of LLPS implicated in cancer are responsible for gene expression-related processes, as well as controlling the creation, breakdown and localization of proteins (**Figure 3**). While these processes are often modulated in cancer in general, LLPS-related proteins play a disproportionately large role in their modulation. The unique nature of cancer-associated LLPS scaffolds becomes even more evident when moving to a higher level. By defining toolkits and supertoolkits, i.e. higher and higher level aggregates of GO terms, it becomes clear that LLPS scaffolds are primarily centered on the maintenance and organization of the genetic material and the regulation of protein availability, as opposed to response to stimuli and the flow of information inside the cell, which are most characteristic of cancer drivers in general (**Figure 3**). At a higher functional level, all these cancer-related processes translate into cellular phenotypes, often referred to as the ten hallmarks of cancer^52^. In this regard, the observed aggressive tumorigenic property can be attributed to the fact that all hallmarks of cancer can emerge from the mutation of LLPS scaffolds, and most hallmarks are significantly enriched in these proteins (**Figure 2**). Furthermore, cancer-driving LLPS scaffolds are often multifunctional proteins in the hallmark space, thus their mutations can contribute to several hallmarks at once, driving tumorigenesis and cancer progression more efficiently.

The aggressive dominant cellular effect of fusions created by LLPS scaffolds is further exacerbated by the fact that the resulting OFPs tend to be modular, large and largely disordered^53^, meaning that finding a single compound to inhibit them is likely to be challenging. In reality, none of the cancer-driving LLPS scaffolds in our dataset has any FDA-approved drug (**Figure 4A**), in line with previous studies of LLPS-prone OFPs^64^. When targeting LLPS-prone proteins or OFPs, many factors need to be considered, for instance, that the partitioning, concentration and activity of cancer drugs may be influenced by the physicochemical attributes of the MLOs^108^. Despite these difficulties, there are a few drugs under development that could target a limited set of LLPS-prone fusion proteins, such as the BRD4-NUTM fusion in midline carcinoma^87^ or LLPS scaffolds fused to RTKs or other kinases potentially being amenable for treatment with kinase inhibitors^55,72,107^. Recently, Wang *et al.* set up a high-throughput imaging-based assay (DropScan) to reassess anticancer drugs as condensate inhibitors, and managed to identify a handful of compounds of low target-specificity that efficiently dissolved condensates of transcriptional OFPs, further validating the direct condensate modulation approach^51^. Furthermore, the pathogenic effects of certain LLPS-prone fusions could potentially be targeted indirectly, through modulating, for instance, their crucial interaction partners, transcriptional targets or the activity of certain chromatin remodeling complexes^64,85,109,110^.

Finding novel strategies for targeting LLPS-inducing OFPs is not just a matter of combating a few obscure cancer cases. While our current analysis only encompasses 69 such fusions due to the limited number of experimentally validated LLPS scaffolds, the true number of OFPs with LLPS scaffolding properties is likely to be much higher, as also suggested by Tripathi *et al.*^50^. We observed a general increase in associations between LLPS-related functions that increase the valency and the interaction capacity of the generated OFPs, such as oligomerization, protein domain specific binding, RNA binding and DNA binding (**Figure 6A** upper matrix). Also, through predictions, we found that OFPs in general display a very high propensity for LLPS, way higher than cancer drivers in general, and strikingly, on par with experimentally validated LLPS scaffolds (**Figure 8**). This is likely due to cases where the individual constituent proteins of the fusion construct cannot induce phase separation, but the fusion protein can, such as the EML4-ALK and CCDC6-RET fusions^76,111,112^. In this light, finding the currently missing drugs to shut down OFPs^64^, to disrupt the condensation enabled by them^51,113^, and to offset their downstream effects^64,85,109^ could provide cancer drugs widely applicable to diverse cancer incidences previously defying standard treatments.

**Figure 8:**
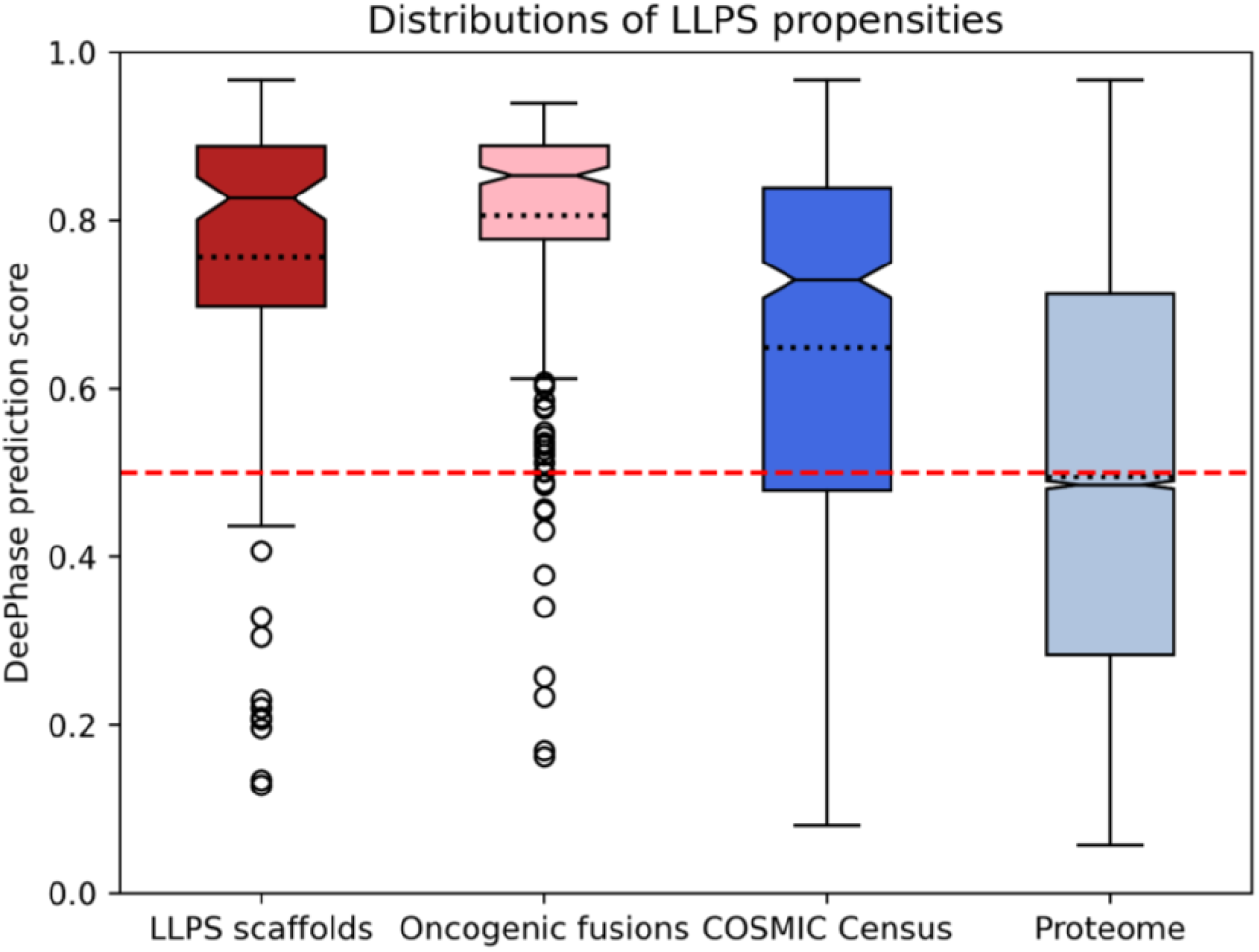
Distributions of predicted LLPS propensities for four different protein groups. LLPS propensity was predicted by DeepPhase for 4 groups of proteins: LLPS scaffolds (red), oncogenic fusion proteins (pink), cancer drivers from COSMIC Census (dark blue) and the whole human proteome (light blue). The continuous horizontal line on the boxes shows the median, while the dotted line indicates the mean of the distribution. The red dashed line is drawn at 0.5, the cutoff value for LLPS on the DeePhase score.

## Data and methods

### 1. Assembly of datasets

### 1.1. LLPS-related proteins

LLPS scaffold proteins were taken as the consolidated dataset of 87 proteins from^7^, and were extended with 54 manually curated proteins totalling 141 high-confidence human LLPS scaffolds. 344 human LLPS regulators were derived from DrLLPS^19^ and 3,503 clients from various data resources (PhaSepDB v1.3^20^, DrLLPS^19^, MSGP^114^, RNP granule DB^115^, MiCroKiTS^116^) that provide information on the localizations of proteins to MLOs (see **Table S1**). Additionally, another larger set of less confident LLPS scaffolds (containing a certain number of clients that have been demonstrated to partition into already existing condensates in *in vitro* experiments) were retrieved from a newer version of PhaSepDB (version 2.1)^117^ presenting a dataset of 859 proteins from low throughput experiments. The obtained proteins were filtered for unique human proteins and the resulting set of 271 proteins (**Table S4**) (which contains 56 COSMIC Census proteins) was used as an independent alternative of our high-confidence scaffold dataset.

### 1.2. Cancer drivers from the COSMIC Census and their actionability from OncoKB

Somatic cancer driver proteins were taken from the Census of COSMIC v95 (**Table S6**). Both tier 1 and tier 2 proteins were used, together with annotations of dominant mutation types, involvement in cancer hallmarks, and molecular roles. Actionability data was taken from OncoKB^118^. Only proteins with an actionability level 1 or 2, i.e. proteins for which there exists at least one FDA-approved drug (level 1), or a drug that is used as standard care (level 2) were considered actionable in our analyses.

### 1.3. Germline cancer-related proteins

Germline cancer-related genes with mutations in protein coding regions were obtained from ClinVar^119^ and HUMSAVAR (https://www.uniprot.org/docs/humsavar) (both downloaded in September 2021) and limited to records with the cancer-related disease terms available in **Supplementary data file 1**. ClinVar classifies phenotype-genotype relationships into groups ranging from less reliable to definite links. To achieve high confidence, entries with “limited”, “disputed”, “no known disease relationship”, “refuted” relationships were not accepted. HUMSAVAR genotype-phenotype records were filtered for the ones accompanied by OMIM phenotype identifiers. The resulting dataset of proteins is available as **Table S2**.

### 1.4. Neurodegenerative disease-linked proteins

ClinVar^119^ and HUMSAVAR (https://www.uniprot.org/docs/humsavar) were downloaded in September 2021, and genes that have neurodegenerative disease-linked mutations in protein coding regions were selected. To achieve this, the dataset was filtered for a curated list of expressions related to neurodegenerative diseases – the precise search terms are available in **Supplementary data file 1**, while the resulting dataset is available as **Table S2**. Entries with the four least confident phenotype-genotype relationship categories were excluded again, as explained previously in *Data and methods section 1.3*. Finally, the remaining mutations that did not match either germline cancer-or neurodegenerative disease-associated terms were included into the “other hereditary diseases” category.

### 1.5. Amyloid fiber-forming proteins

Human amyloid fiber-forming proteins were retrieved from the AmyPro database^120^ in August 2021 (**Table S5**).

### 1.6 Subcellular localization for the full human proteome

We defined the subcellular localization for each protein in the human proteome by integrating data from Gene Ontology annotations in UniProt (GOA), UniProt annotations, the Human Transmembrane Proteome (HTP)^121^, MatrixDB^122^, and MatrisomeDB^123^. We divided the UniProt and the Gene Ontology annotations (GOA) into tier 1 (more reliable) and tier 2 (less reliable) annotations, depending on the attached evidence codes. For UniProt, annotations with the evidence codes ECO:0000269 or ECO:0000305 are considered as tier 1, while annotations with evidence codes ECO:0000250, ECO:0000255, or ECO:0000303 are tier 2. For Gene Ontology, annotations with evidence codes IDA, IMP, IPI, IGI, EXP, IBA, IKR, TAS, NAS, IC, or ND are tier 1, while annotations with evidence codes HDA, ISS, ISA, RCA, ISO, ISM, IGC, or IEA are tier 2.

Based on these, each protein was assigned exactly one broad localization. It was considered to be a transmembrane protein (TMP), if it is assigned the ‘integral component of membrane (GO:0016021)’ GO term in tier 1 GOA annotations, or it is annotated as a TMP in HTP with a confidence score over 85, or is annotated in HTP as a TMP with a confidence score above 50 and is also annotated as a TMP in GOA (either tier). TMPs were further categorized into *Plasmamembrane TMPs* (if they had the ‘plasma membrane (GO:0005886)’ GO annotation in either tier of GOA, or had any of the following terms in their tier 1 or tier 2 UniProt annotations: cell membrane, postsynaptic density membrane, flagellum membrane, cilium membrane, dendritic spine membrane, filopodium membrane, growth cone membrane, invadopodium membrane, lamellipodium membrane, microvillus membrane, podosome membrane, pseudopodium membrane, ruffle membrane, stereocilium membrane), *Internal membrane TMPs* (if annotated with any of the intracellular localizations, see below), *External membrane TMPs* (if annotated with any of the extracellular localizations, see below), and *TMPs in unknown membrane* (if none of the previous categories could be assigned).

Proteins (TMP and non-TMP) were annotated with the following intracellular localizations:

● nuclear, if it has the ‘nucleus (GO:0005634)’ term attached in GOA tier 1, or the ‘Nucleus’ term attached in UniProt tier 1; or if it has no tier 1 annotations, but is attached the same GO term in GOA tier 2, or the same UniProt term in tier 2;
● cytosol, if it has the ‘cytosol (GO:0005829)’ term attached in GOA tier 1, or the ‘Cytosol’ term attached in UniProt tier 1; or if it has no tier 1 annotations, but is attached the same GO term in GOA tier 2, or the same UniProt term in tier 2;
● nucleus/cytoplasm shuttling, if it can be annotated both as a nuclear and a cytosolic protein based on the above definitions;
● ER, if it has the ‘endoplasmic reticulum (GO:0005783)’ term attached in GOA tier 1, or any of the ‘Endoplasmic reticulum’, ‘Endoplasmic reticulum lumen’ and ‘Endoplasmic reticulum membrane’ terms attached in UniProt tier 1; or if it has no tier 1 annotations, but is attached the same GO term in GOA tier 2, or the same UniProt term in tier 2;
● Golgi, if it has the ‘Golgi apparatus (GO:0005794)’ term attached in GOA tier 1, or if it has no tier 1 annotations, but is attached the same GO term in GOA tier 2, or the ‘Golgi apparatus’ annotation in UniProt tier 2;
● cytoskeleton, if it has the ‘cytoskeleton (GO:0005856)’ term attached in GOA tier 1, or the ‘cytoskeleton’ term attached in UniProt tier 1; or if it has no tier 1 annotations, but is attached the same GO term in GOA tier 2, or the same UniProt term in tier 2;
● mitochondrium, if it has the ‘mitochondrion (GO:0005739)’ term attached in GOA tier 1, or the ‘Mitochondrion’ term attached in UniProt tier 1; or if it has no tier 1 annotations, but is attached the same GO term in GOA tier 2, or the same UniProt term in tier 2;
● other intracellular organelle, if it has the ‘intracellular anatomical structure (GO:0005622)’ term attached in GOA tier 1, or the ‘Cytoplasm’ term attached in UniProt tier 1, and cannot be assigned any of the above, more specific localizations; or if it has no tier 1 annotations, but is attached the same GO term in GOA tier 2, or the same UniProt term in tier 2.

Or if none of these could be assigned, then one of the following extracellular localizations:

● extracellular vesicle, if it has any of the ‘exosome (GO:0070062)’, ‘microvesicle (GO:1990742)’, or ‘prominosome (GO:0071914)’ terms attached in GOA tier 1, or either the ‘extracellular vesicle’ or ‘extracellular exosome’ term attached in UniProt tier 1; or if it has no tier 1 annotations, but is attached the same GO term in GOA tier 2, or the same UniProt term in tier 2;
● extracellular, if it has any of the ‘extracellular space (GO:0005615)’, ‘collagen trimer (GO:0005581)’ or ‘complex of collagen trimers (GO:0098644)’ terms attached in GOA tier 1, or the ‘Mitochondrion’ term attached in UniProt tier 1; or if it has no tier 1 annotations, but is attached the same GO term in GOA tier 2, or the same UniProt term in tier 2.

If none of these terms could be defined for the protein, it was classified as ‘Unknown localization’. The list of subcellular localizations for all human proteins is shown in **Table S3**.

### 1.7. Randomized selections of human proteins from UniProt to gain unbiased background sets for statistical comparisons

Cancer-associated proteins in the COSMIC Census are usually highly researched owing to their established disease link. Thus, these proteins usually have high annotation scores in UniProt with many Gene Ontology (GO) terms associated with them. In addition, cancer-associated proteins are often plasmamembrane receptors and transcription factors, leading to a non-uniform distribution across various subcellular localizations. These deviations from the average human proteins would lead to severe annotation biases in our enrichment calculations. Therefore, in each such analysis, we compare the COSMIC Census proteins to random sets of proteins that share the same annotation score in UniProt (5 out of 5) and the same distribution across various subcellular localizations (as defined above). We generated 1000 sets of randomly selected proteins from the human proteome that have the same number of proteins, all with a UniProt annotation score of 5, and the same subcellular localization distribution as the set of proteins we are assessing, for instance, COSMIC Census proteins. This procedure was applied to gain unbiased randomized background sets for all disease protein sets, separately (see **Supplementary data files 2-5**), as well as for oncogenes and tumor suppressors (**Supplementary data files 6-7**).

### 1.8. Oncogenic fusions

#### 1.8.1. Assembling data on the oncogenic fusion proteins of all COSMIC Census proteins

We performed a comprehensive, protein-level manual curation of the OFPs of all COSMIC Census proteins, where fusion was provided as a dominant mutation type. For these proteins, each of their fusion partners listed either by COSMIC or UniProt were collected, and the corresponding fusion gene pairs annotated, totaling 450 unique gene pairs.

Fusions of the same two genes that only differ in their fusion breakpoints were considered as variants of the same fusion and not as distinct fusions, so only one representative was annotated. Only in-frame fusions of two different genes were considered, where the resulting fusion protein contained in-frame portions of any size of both encoded proteins, including those cases where a short non-coding (usually intronic) segment separates two in-frame protein-coding regions. For the fusion gene pairs for which at least one COSMIC sample and fusion identifier was available, we selected the most frequently occurring fusion setup/breakpoint (the one indicated with the most samples) and made an attempt to annotate the exact protein boundaries based on that. A preference was given to fusion identifiers/transcript-level descriptions, wherein the transcript boundaries were well-defined (lacking +/- and characters marking lack of confidence in fusion boundaries). The UniProt isoforms corresponding to the indicated Ensemble transcript identifiers could be unambiguously identified in each case. For the N-terminal fusion partner we obtained the protein boundary by taking the indicated fusion breakpoint of the transcript, subtracting the length of the 5’UTR and dividing the remaining number (length of CDS before the breakpoint) by 3. The integer portion of the result was accepted as the protein level fusion breakpoint for the N-terminal gene. If the remainder after dividing by 3 was not zero, we made a note of that to see if the fusion is in-frame or not. For the transcript of the C-terminal gene, based on the first nucleotide indicated by COSMIC to be part of the fusion, we calculated the length of the CDS that was missing from the fusion and divided it by 3 to see how many residues were missing from the N-terminus of the protein. The remainder (if any) was compared to the previously noted remainder of the N-terminal gene. If the two remainders were equal, that means that the N-terminal portion of the first gene could substitute for the N-terminal portion of the second gene in a way that the reading frame was preserved, so the fusion was accepted to be an in-frame fusion.

The fused regions’ boundaries were annotated in a way that only residues entirely encoded by the respective coding regions have been accepted. De novo residues made up by the fused codons or originally non-coding spacer regions were not added to any of the two protein regions, but were noted as middle residues (if they could be identified). In the majority of the cases, where the fusions were annotated based on COSMIC fusion transcripts, the middle residues could not be identified, only the number of nucleotides the two different genes contributed to the encoding of the middle residue. We noted this as 2+1 or 1+2, where the contribution of the N-terminal gene is the first number and the contribution of the C-terminal gene is the second number.

For the fusion partners, where the fusion breakpoints were not provided by COSMIC or UniProt, those were obtained through comprehensive literature curation of the associated articles. In these cases, we revisited the original articles where the precise boundary of the fusion was described (usually the first article reporting on the fusion of the two genes).

The integrated table with the resulting manually curated oncogenic fusions from UniProt and COSMIC is available as **Table S11**. In the table it is also provided if at least one of the fusion partners of the OFPs is an LLPS driver, so those could be separately analyzed.

## 2. Assembly of the molecular toolkits

Inspired by an earlier cancer analysis paper^124^, we created a large compilation of 21 molecular toolkits belonging to 5 higher level categories (‘supertoolkits’), defined based on GO annotations available for proteins^125,126^. These encompass a diverse set of functions that cover a broad range of actions proteins carry out within human tissues comprising amongst others genome organization, regulation of protein availability, transport-related, signaling-related and other processes (**Figure 3A**).

Description of the toolkit and supertoolkit definitions along with the exact GO terms defining the given molecular toolkits are listed in **Tables S8 and S9**.

Toolkit enrichment for LLPS categories (scaffold, regulator, client) was compared to the presence of toolkit terms in cancer drivers, while the enrichment of toolkits for cancer drivers was contrasted with an equivalent, unbiased, randomized background (**Supplementary data file 2**). Both were quantified by fold enrichment values and the p-values of Fisher’s exact tests (**Table S10**).

## 3. Domain mapping, functional annotation of domains and enrichment analysis

### 3.1. Identifying InterPro, Pfam, and UniProt regions

Pfam^127^ (downloaded on February 24, 2023) and InterPro^128^ (version 5.56) annotations were used to scan both cancer driver proteins, as well as OFPs. UniProt region annotations were taken from the UniProt database, downloaded on October 7, 2022. UniProt regions were assigned to OFPs if the fusion construct contained at least 10% of the residues in the original region. While this cutoff is low, for structured regions (such as domains, DNA-binding regions, etc), it is virtually always the case that the fusion product either contains all of the region or none of it^53^. Fractions of UniProt regions only show up in fusion constructs where the region denotes a region without well-defined tertiary structure or if the region is repetitive in nature, such as coiled coils. In these cases, the permissive 10% cutoff ensures that we capture functionality arising from only a portion of the region.

### 3.2. Gene Ontology annotations of Pfam, InterPro, and UniProt regions

Identified InterPro and Pfam regions were attached with Gene Ontology (GO) terms using various sources (**Tables S12 and S13**). The InterPro2GO and Pfam2GO mappings were downloaded from the EBI servers on June 20, 2022. In addition, we further annotated various Pfam regions based on literature sources: RNA binding function (GO:0003723) was assigned to Pfam domains and families in the EuRBPDB^129^, while DNA binding function (GO:0003677) based on prior efforts of Malhotra & Sowdhamini^130^, and phospholipid binding (GO:0005543) based on MBPpred^131^. Protein dimerization (GO:0046983), homooligomerization (GO:0051260) and complex oligomerization (GO:0051259) were annotated based on relevant articles^132^ and additional manual curation efforts. Chromatin modifiers were stringently recurated starting from an earlier study^133^, and domains were functionally mapped to a small set of GO terms with varying annotation depth related to chromatin modification.

For UniProt regions/domains/sites/motifs annotated to more than one oncogenic fusions, these protein components were functionally characterized by GO terms using manual curation (**Table S14**). Additionally, some of the UniProt regions occurring only once in the protein set were also functionally annotated. In total more than 800 GO terms were manually assigned to these UniProt regions from a set of 27 GO-defined molecular functions or biological processes. One of the most common UniProt regions in our set was the term “coiled-coil region” (126 occurrences), for which functional annotation is less trivial. For simplicity, the GO term “protein homooligomerization” (GO:0051260) was assigned to it as a proxy.

InterPro and Pfam domains listed in the ELM database^134^ (downloaded on September 28, 2022) as binding partners for any motif were annotated with the ‘GO:0019904, protein domain specific binding’ term.

The InterPro/Pfam/UniProt - GO associations were then used to attach GO annotations to cancer proteins (**Table S15**) and OFPs (**Table S16**). These GO terms were then mapped to a GO subset (GO Slim) that represents biologically relevant, fairly specific yet high level processes and functions (**Table S17**).

### 3.3 Mapping the minimal regions of LLPS scaffolds and labeling them by annotations

The minimal LLPS regions of the fusion-forming LLPS scaffolds were derived from *in vitro* experiments describing the minimal requirements of LLPS in the references provided in **Table S21**. Annotation of the term “driving biological condensation” to (fusion) proteins followed similar rules as for UniProt annotations. This label was considered together with the GO terms and is also shown in **Tables S15-20**.

### 3.4. Enrichment analysis of functional region associations

We analyzed how commonly pairs of functional categories (defined by GO terms) of different protein domains and regions associate to each other in cancer proteins in general, and also in oncogenic fusion proteins in specific (**Tables S18-19**). Functional term associations were defined by the overlap coefficient metric (also known as Szymkiewicz–Simpson coefficient): OC(X,Y) = |X∩Y| / min(|X|,|Y|).

Enrichment of associations was defined as a simple difference of overlap coefficients (ΔOC = OC1 - OC2). **Table S20** shows the differences of these overlap coefficients between the cancer proteins and OFPs.

## 4. Large-scale prediction of LLPS propensity

DeePhase^135^, Droppler^136^, PSPredictor^137^ and GraPES^138^ were benchmarked on our LLPS scaffold set. GraPES predictions were accessed from the online database, and the MaGS Z-scores were converted using the (z+1)/3 formula and capped at [0,1]. The rest of the predictors gave numbers within [0,1]. While DeePhase and PSPredictor are parameter-free and only take the sequence as input, Doppler had to be run by setting the experimental conditions for which we chose the default parameters (T=37°C, c=10uM, pH=7, I=0.01M, crowder=None).

The LLPS propensity scores for each protein were compiled as distributions for each predictor, and evaluated as quartiles and upper/lower interquartile ranges * 1.5 (IQR*1.5) visualized as box-whisker plots (python 3.10; using package ‘matplotlib’ version 2.6.3). DeePhase was selected for proteome-wide LLPS propensity prediction as its propensity distribution including the data points ranging from the upper to the lower whisker were best overlapping with the 0.5-1.0 normalized LLPS propensity range, while other predictors more often predicted lower LLPS propensities (<0.5) for experimentally validated LLPS scaffolds. Proteome-wide LLPS propensity prediction (**Table S22**) was performed on the UniProt-assembled human reference proteome (UP000005640) downloaded in May 2022 (**Table S3**). From this dataset two selections were made: a subset for the LLPS scaffolds and a subset for the cancer drivers from COSMIC Census (**Supplementary data file 8**). Lastly, LLPS propensities were also predicted (**Table S23**) by DeePhase for reconstituted sequences of oncogenic fusion proteins (*see Data and methods subsection 1.8* and **Supplementary data file 9**).

## 5. Statistical analyses

χ2 statistics were applied to address the statistical significance of overlaps between LLPS scaffolds and various disease-associated proteins using the reviewed human proteome (20,359 proteins) from UniProt as a background. Generally human LLPS proteins are very well annotated, in comparison to many non-LLPS proteins. Therefore, to obtain a proper baseline devoid of any bias resulting from retrieving random proteins from human proteome which are either understudied or belonging to a subcellular localization irrelevant to LLPS, for example extracellular proteins, we filter the whole human proteome to proteins that are similarly well annotated and distributed within the cell. Subsequently, the obtained subset was used as a background for performing the random selections of proteins to serve as reference sets for the significance tests.

Due to the smaller data sizes, we chose to apply Fisher’s exact test using the 713 COSMIC cancer genes as background in statistical analyses of the association between LLPS and cancer hallmarks or other characteristics including molecular toolkits. In the case of molecular toolkits, the overrepresentation of cancer drivers among the proteins of the human proteome belonging to each toolkit was evaluated by comparisons to a background consisting of 100 randomized datasets with an equivalent number of proteins. The overrepresentation values of cancer drivers served as a baseline to evaluate the extent and significance of toolkit enrichments of the cancer-associated proteins of the 3 LLPS groups.

## Acknowledgements and funding

This project has been implemented with the support provided by the Ministry of Innovation and Technology of Hungary from the National Research, Development and Innovation Fund, financed under the K-124670 and K-131702 funding schemes granted to P.T. and the FK-128133 and FK-142285 funding schemes granted to R.P. R.P is a holder of the János Bolyai Research Fellowship of the Hungarian Academy of Sciences (BO/00174/22). This work was supported by an EC H2020-WIDESPREAD-2020-5 Twinning grant ‘PhasAge’ (no. 952334 to P.T.). N.F is a PhD fellow supported by an FWO fellowship in fundamental research (FWOTM1124). T.L. is a postdoctoral innovation mandate holder (HBC.2022.0194) of the Flanders Innovation & Entrepreneurship Agency (VLAIO). B.M. thanks ALSAC for funding and support for his research.

