## Supplementary figures and information for "Phase-separating fusion proteins drive cancer by dysregulating transcription through ectopic condensates"

[#] co-first authors
* corresponding authors

### Supplementary figures

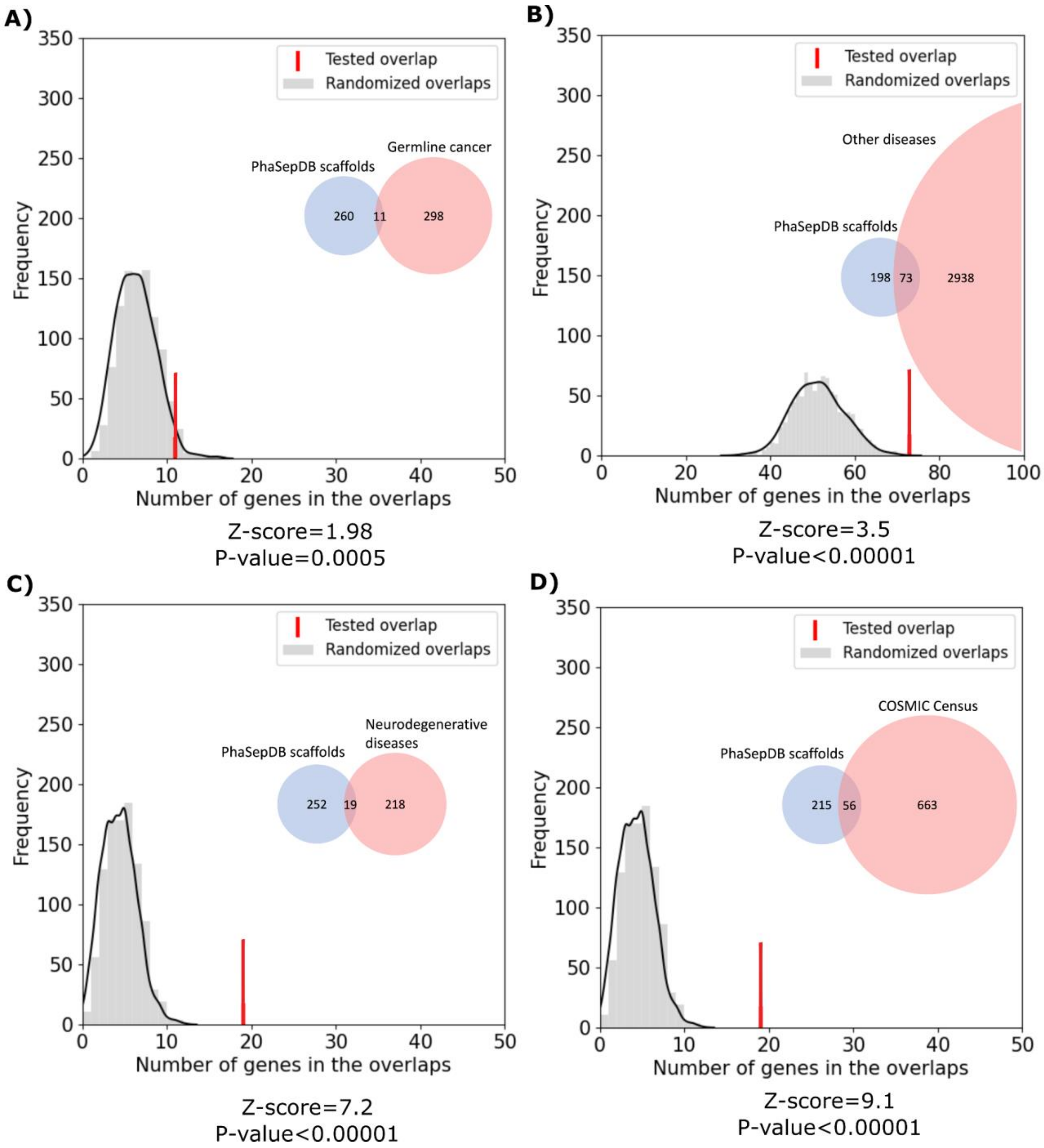


**Figure S1: Overlap between PhaSepDB-derived LLPS scaffolds and various disease-associated proteins.** Gray distributions show the expected overlap between PhaSepDB LLPS scaffolds and the four classes of disease-associated proteins: (**A**) germline cancer, (**B**) other human diseases, (**C**) neurodegenerative diseases and (**D**) somatic cancer. Distributions were calculated from 1000 random generated sets of human proteins with subcellular localizations and levels of annotation matched to the real disease protein sets (*see Data and methods*). Red bars mark the observed overlap between the true protein sets with the corresponding Z-scores and p-values indicated below the graphs.

Inset Venn diagrams show the number of proteins in each set and the observed overlap between them with circle areas being proportionate to the corresponding set sizes.

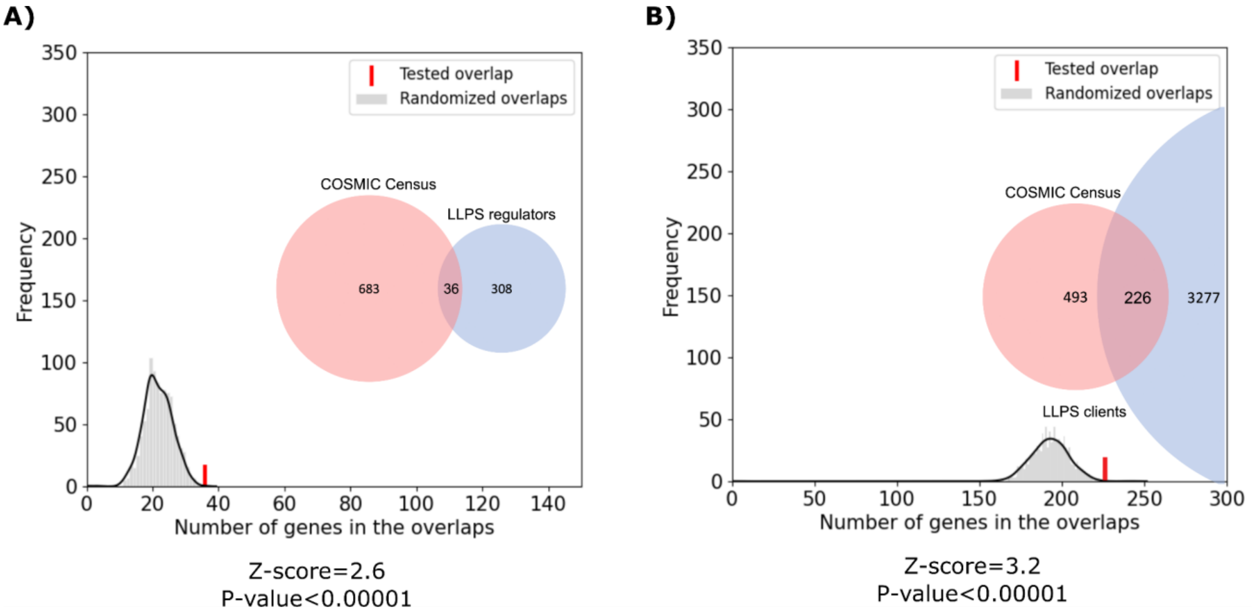


**Figure S2: LLPS regulators and clients are significantly overrepresented among COSMIC Census cancer driver proteins, but more moderately than LLPS scaffolds.** Gray distributions show the expected overlap between (**A**) LLPS regulators and (**B**) clients with COSMIC Census proteins. Distributions were calculated from 1000 random generated sets of human proteins with subcellular localizations and levels of annotation matched to the COSMIC Census (*see Data and methods*). Red bars mark the observed overlap between the true protein sets with the corresponding Z-scores and p-values indicated below the graphs. Inset Venn diagrams show the number of proteins in each set and the observed overlap between them with circle areas being proportionate to the corresponding set sizes.

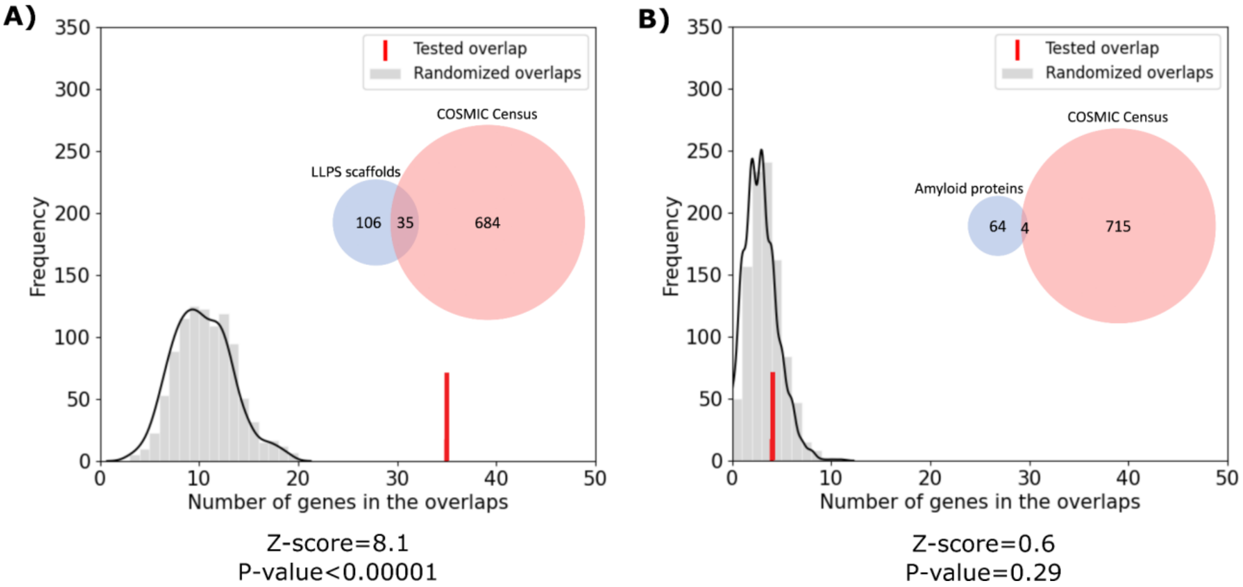


**Figure S3: LLPS scaffolds show larger overlap with somatic cancer driver proteins than amyloid driver proteins.** Gray distributions show the expected overlap between human (**A**) LLPS scaffolds and (**B**) amyloid-forming proteins with COSMIC Census proteins. Distributions were calculated from 1000 random generated sets of human proteins with subcellular localizations and levels of annotation matched to the COSMIC Census (*see Data and methods*). Red bars mark the observed overlap between the true protein sets with the corresponding Z-scores and p-values indicated below the graphs. Inset Venn diagrams show the number of proteins in each set and the observed overlap between them with circle areas being proportionate to the corresponding set sizes.

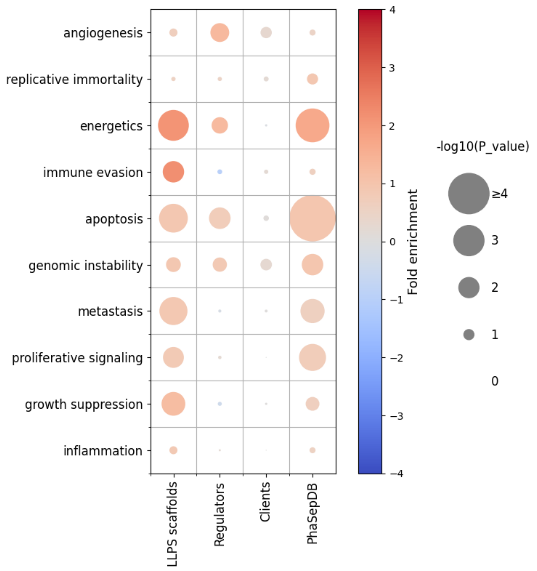


**Figure S4: Association of LLPS-related proteins (self-curated data and PhaSepDB scaffold dataset) with cancer hallmarks.** The color of the circles in the heatmap represents the fold enrichment, while the size represents the significance of overrepresentation/depletion for our three groups of LLPS-related proteins and the PhaSepDB-derived scaffolds in the ten hallmarks of cancer.

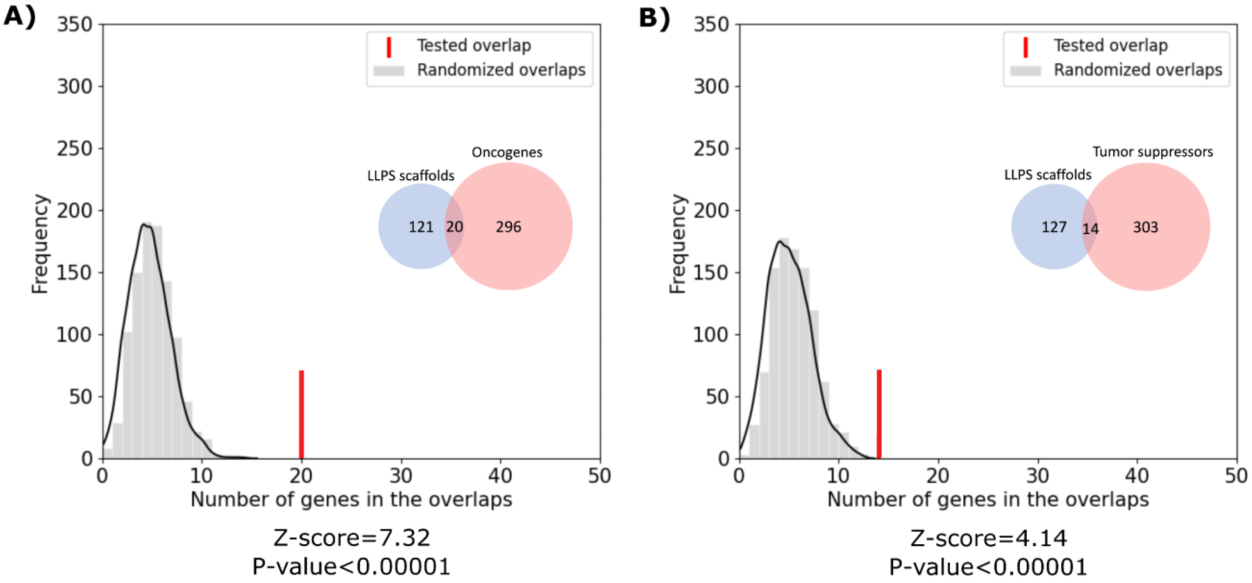


**Figure S5: LLPS scaffolds are significantly overrepresented among both oncogenes and tumor suppressors, but more prominently among oncogenes.** Gray distributions show the expected overlap between LLPS scaffolds and (**A**) oncogenes, or (**B**) tumor suppressors. Distributions were calculated from 1000 random generated sets of human proteins with subcellular localizations and levels of annotation matched to the real (**A**) oncogene, (**B**) tumor suppressor protein sets (*see Data and methods*). Red bars mark the observed overlap between the true protein sets with the corresponding Z-scores and p-values indicated below the graphs. Inset Venn diagrams show the number of proteins in each set and the observed overlap between them with circle areas being proportionate to the corresponding set sizes.

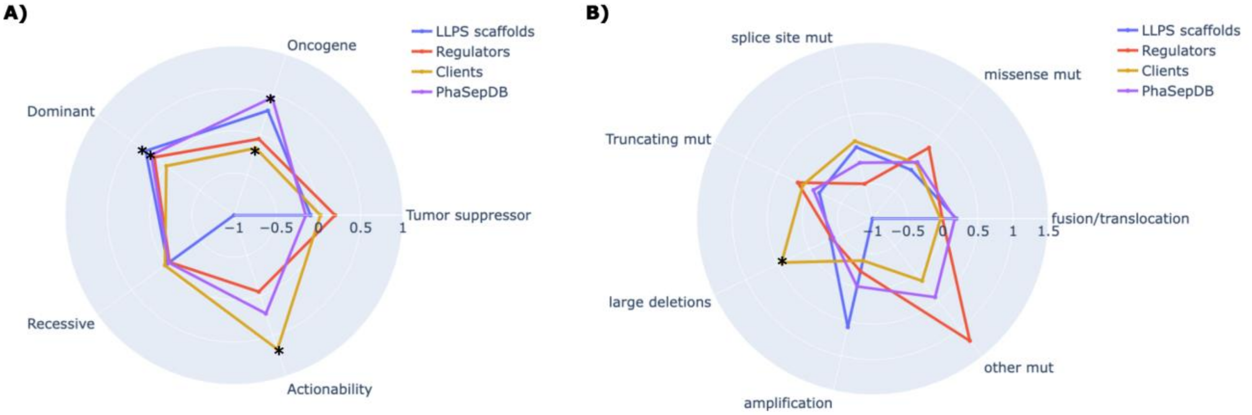


**Figure S6: Characteristic features of cancer-associated LLPS proteins from our own dataset (LLPS scaffolds - blue, LLPS regulators - red, LLPS clients - yellow) and from PhaSepDB (purple).** (**A**) The radar chart plots the fold enrichments of the four classes of cancer-associated LLPS proteins in cancer drivers that are oncogenes or tumor suppressors; that are affected by dominant or recessive mutations; and those with available FDA-approved drugs (actionability). **(B)** Fold enrichments of the four classes of cancer-associated LLPS proteins in cancer drivers with different dominant mutation types.

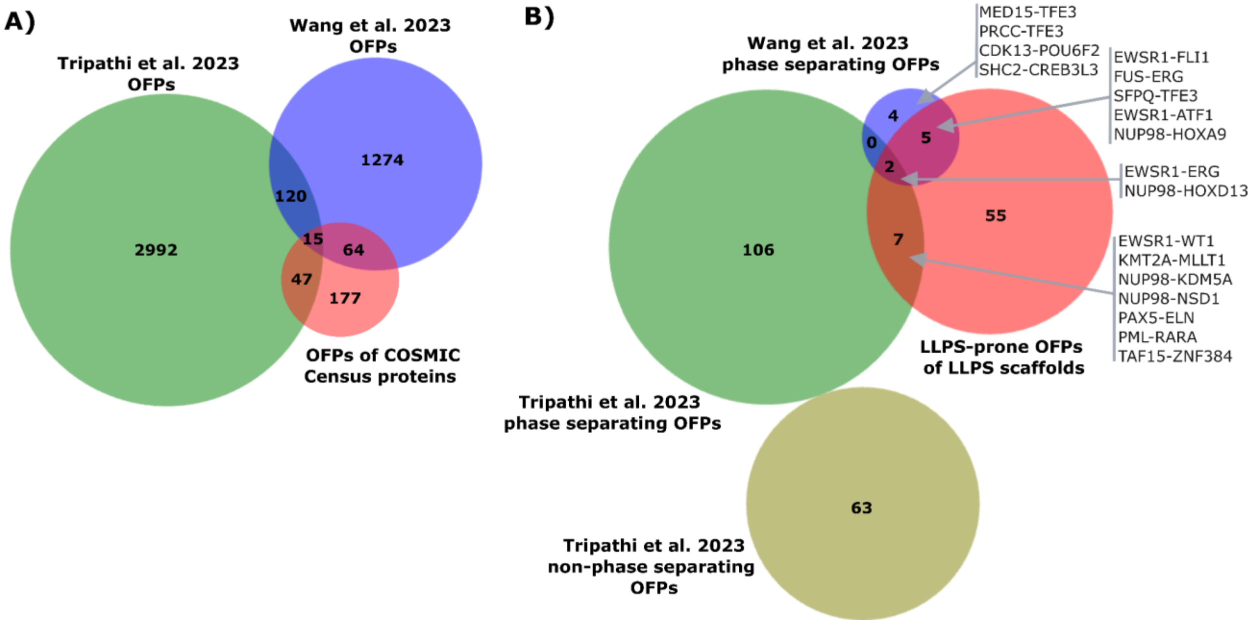


**Figure S7: Overlap of oncogenic fusion protein (OFP) datasets derived from various sources.** (**A**) Overlap of our COSMIC-derived OFP dataset with the OFP datasets analyzed by Tripathi *et al.*[50] and Wang *et al.*[51]. (**B**) Overlap of our LLPS-prone OFPs dataset of the curated LLPS scaffolds with the OFPs displaying punctate cellular localization in HeLa cells by Tripathi *et al.*, the few OFPs demonstrated to undergo LLPS *in vitro* and *in cellulo* by Wang *et al.*[51], and with those showing a diffuse cellular localization in HeLa cells (therefore probably failing to undergo LLPS) by Tripathi *et al.*[50]. The Venn diagrams reveal poor overlaps both between the entire sets of analyzed OFPs and the subsets of OFPs tested/annotated for driving LLPS.

#### Supplementary tables

**File: Supplementary tables S1-S7.xlsx**

**Table S1:** LLPS-associated proteins with roles and data sources indicated.
**Table S2:** Disease-associated proteins with disease category
**Table S3:** Subcellular localizations and annotation scores for all human proteins
**Table S4:** Human LLPS scaffolds obtained from PhaSepDB 2.1 with involvement in cancer indicated
**Table S5:** Amyloid forming human proteins from the AmyPro database
**Table S6:** List of COSMIC Census cancer drivers with annotations
**Table S7:** Enrichments calculated for COSMIC annotated features

**File: Supplementary tables S8-S10.xlsx**

**Table S8:** Toolkit definitions and descriptions
**Table S9:** Supertoolkit definitions and descriptions
**Table S10:** Enrichments calculated for the toolkits

**File: Supplementary tables S11-S21.xlsx**

**Table S11:** List of collected OFPs with boundaries and annotations
**Table S12:** GO annotations assigned to Pfam domains
**Table S13:** GO annotations assigned to InterPro domains
**Table S14:** GO annotations assigned to UniProt regions
**Table S15:** GO annotations for cancer proteins
**Table S16:** GO annotations for OFPs
**Table S17:** Term definitions of applied GO Slim, number of cancer proteins/OFPs belonging to each term
**Table S18:** Overlap coefficients between the occurrences of pairs of GO terms in cancer proteins
**Table S19:** Overlap coefficients between the occurrences of pairs of GO terms in OFPs
**Table S20:** Overlap coefficient differences between the occurrences of pairs of GO terms in cancer proteins and OFPs
**Table S21:** Minimal LLPS driver region boundaries for the cancer fusion-associated LLPS scaffolds

**File: Supplementary table S22-S23.xlsx**

**Table S22:** LLPS propensity prediction by DeePhase for the human proteome
**Table S23**: LLPS propensity prediction by DeePhase for oncogenic fusions of cancer driver proteins

#### Supplementary data files

**Supplementary data file 1:** List of terms used for filtering germline cancers and neurodegenerative diseases

**Supplementary data file 2:** Random protein selections from UniProt for background calculations for somatic cancer proteins (COSMIC Census).

**Supplementary data file 3:** Random protein selections from UniProt for background calculations for proteins implicated in neurodegenerative diseases.

**Supplementary data file 4:** Random protein selections from UniProt for background calculations for proteins implicated in other diseases.

**Supplementary data file 5:** Random protein selections from UniProt for background calculations for proteins implicated in germline cancers.

**Supplementary data file 6:** Random protein selections from UniProt for background calculations for oncogenes

**Supplementary data file 7:** Random protein selections from UniProt for background calculations for tumor suppressors

**Supplementary data file 8:** COSMIC Census driver sequences

**Supplementary data file 9:** Reconstituted OFP sequences
